# Mitochondrial sites of contact with the nucleus aid in chemotherapy evasion of glioblastoma cells

**DOI:** 10.1101/2024.08.27.608373

**Authors:** Daniela Strobbe, Mardja Bueno, Claudia De Vitis, Sarah Hassan, Danilo Faccenda, Krenare Bruqi, Elena Romano, Lucia Pedace, Andrey A. Yurchenko, Gurtej K Dhoot, Ivi J Bistrot, Fabio Klamt, Luana S. Lenz, Eduardo Cremonese Filippi–Chiela, Pietro Ivo D’Urso, Imogen Lally, Eveline Miele, Laura Falasca, Sergey Nikolaev, Rita Mancini, Federico Roncaroli, Guido Lenz, Michelangelo Campanella

## Abstract

Glioblastoma (GBM) is the most common form of a malignant primary brain tumour in adults for which therapeutic options are minimal. The rapid onset of the resistance mechanisms against the chemotherapeutic agent Temozolomide (TMZ), the first line of pharmacological care for patients, prevents the long-term validity of this approach. The underpinning biology for this remains poorly understood thus compromising the efficacy of this approach. The Translocator Protein (TSPO) is an 18kDa ubiquitous cholesterol-binding molecule on the outer membrane of mitochondria (OMM). Upregulated in cancers TSPO is required to form contacts between mitochondria and the nucleus termed: Nucleus Associated Mitochondria (NAM). In GBM tissues as well as in 2D and 3D cell cultures we assayed patterns of TSPO expression (i), autophagy/mitophagy (ii), transcription factors (iii) and susceptibility to TMZ-induced demise (iv). Confocal and ultrastructural imaging detailed the organization and redistribution of the mitochondrial network (v).

Our findings show that TMZ exploits mitochondria via TSPO to aid the formation of NAM which couples the expression of the nuclear transcription factor Sterol regulatory element-binding transcription factor 1 (SREBP1) and the stabilization of YAP/TAZ.

Pharmacological modulation of TSPO counteracts all the above and re-instates susceptibility to TMZ-induced demise. NAM is therefore proposed as a variable in the engagement and execution of pro-survival mechanisms in GBM thus offering a means to both insight into the pathophysiology of this disease and offer novel therapeutic strategies.

**Key Points:**

- TMZ exploits TSPO to curb mitochondrial quality control in glioblastoma cells.
- TMZ-mediated MRR is associated with the relocation of mitochondria to the nucleus and modulation of transcriptional factors involved in cholesterol metabolism and adaptation to aggressive growth.
- TSPO represents a pharmacological target to revert chemoresistance in glioblastoma cells.

**Importance of the Study:** This study elucidates a mitochondrion-driven mechanism of chemoresistance in human glioblastoma cells, which depends on the mitochondrial translocator protein TSPO. The administration of TSPO ligands restores susceptibility to TMZ by influencing the dynamics of transcriptional factors associated with cholesterol metabolism and mechanical transduction.

## Introduction

Glioblastoma multiforme (GBM) is the most common and aggressive primary brain tumour in adults (Tykocki & Eltayeb, 2018) with an overall median survival between 12 and 15 months (Michaelsen et al., 2013) and common recurrence (Lawson et al., 2007). The high degree of GBM infiltration in the normal brain, along with organized anatomical structures such as blood vessels and white matter tracts, limits the extent of surgical intervention (Braun & Ahluwalia, 2017).

In addition, the molecular and metabolic heterogeneity of brain tumour cells (Nassiri et al., 2023; Sottoriva et al., 2013) counteract the efficacy of radiotherapy and chemotherapy therapeutic approaches. The alkylating agent temozolomide (TMZ) represents the current standard of care for patients in pre and post-surgical procedures (Colwell et al., 2017; Fekete et al., 2023). TMZ delivers intracellular toxicity targeting nuclear (nDNA) and mitochondrial DNA (mtDNA) (Lee, 2016; Nassiri et al., 2023) by inserting methylene groups at positions N-7 or O-6 of guanine residues.

GBM cells react to genotoxic stress by activating DNA repair enzymes such as the O6-methyl guanine methyl transferase (MGMT) which is upregulated in the majority of glioblastomas (Cho et al., 2023; Oliva et al., 2010). Notably, though, the degree of MGMT expression isn’t the only signature associated with therapy evasion in GBM cells; resistance against TMZ treatment is recorded also in the types of glioblastomas with lower MGMT expression (Zhang et al., 2022). Gaining a deeper understanding of the biological redesign leading to chemoresistance is therefore paramount to learning how the cytotoxic potential of therapeutic approaches could be retained (Aldape et al., 2019).

It is acknowledged that mitochondria partake in the molecular strategies of evasion enacted by cancer cells but their contribution to the GBM cells’ resistance to alkylating agents remains ill-defined (Ammer et al., 2020). In response to stressors, mitochondria contribute to the adaptative response by engaging with the nucleus via a retrograde conduit of communication known as Mitochondrial Retrograde Response (MRR) (Desai et al., 2020; Yang & Kim, 2019). Reprogramming of cancer metabolism and activation of proliferative genes is mediated by MRR which is read by the nuclear accumulation of pro-survival transcription factors (S. Kim & Koh, 2017) such as nuclear factor kappa-light-chain-enhancer of activated B cells (NF-kB) (Quirós et al., 2016).

Recently, our group showed that MRR is facilitated by the formation of contact sites between mitochondria and the nucleus (referred to as Nucleus-associated mitochondria: NAM) (Desai et al., 2020) for which the upregulation of the 18kD Translocator Protein (TSPO), a cholesterol-binding located on the outer mitochondrial membrane (OMM) is required (Strobbe et al., 2021; Teraiya et al., 2023).

TSPO is overexpressed in several types of aggressive tumours (Ammer et al., 2020) including high-grade gliomas for which its theragnostic potential has been proven in pre-operative stratification of patients (Ammer et al., 2020; Carter et al., 2023; Roncaroli et al., 2016) leaving untested the potential in counteracting resistance mechanisms via its targeting. Several are indeed compounds binding to TSPO which have been linked to therapies (Singh et al., 2021). At the cellular level, TSPO is implicated in a broad spectrum of processes encompassing steroid synthesis including the transport of cholesterol to mitochondria (Rupprecht et al., 2010), regulation of proliferation (Wu et al., 2020), oxidative stress (Zeno et al., 2012) and mitochondrial quality control via targeted autophagy (i.e. mitophagy) (Gatliff & Campanella, 2015).

TSPO not only promotes the formation of NAM via the molecules with which it interacts (e.g. ACBD3 and PKA) (Jaipuria et al., 2017) but also aids the redistribution of cholesterol and its oxidized metabolites on the nucleus to positively regulate the transcription of pro-survival genes (Desai et al., 2020).

Cholesterol anabolism and catabolism are closely linked with GBM pathogenicity framed by redesigned energy balance (Pirmoradi et al., 2019). GBM cells engage the YAP and TAZ mediators of the Hippo pathway (hereafter called YAP/TAZ) to promote tissue proliferation and organ growth (Castellan et al., 2021; Sorrentino et al., 2014). Thus YAP/TAZ hold the capacity to reprogram cancer cells into staminal ones to prime tumour initiation, progression, and metastases (Castellan et al., 2021). In addition, they also aid chemoresistance and aggressiveness of the gliomas (Masliantsev et al., 2021). Although direct evidence on the interplay between YAP/TAZ, cholesterol homeostasis and the MVA pathway is currently lacking, a hierarchical link has been previously speculated (Sorrentino et al., 2014) calling for attention to the less investigated mechanisms of cholesterol regulation such as those of mitochondrial origin.

On these premises, we endeavoured to investigate the role of the cholesterol-binding molecule TSPO in the biology of glioblastoma cells to inform a mitochondrial conduit of adaptation to genotoxic stress via cholesterol. Our results show that the chemotherapeutic agent TMZ relies on the retrograde mechanisms of mitochondrial communication with the nucleus to sustain cell death evasion and the contact sites between the two organelles pivot such a response. Above all this study offers novel insights into the role of mitochondria in resistance mechanisms and how this can be overcome by targeting cholesterol.

## Materials and Methods

### Tissue culture

GBM cell lines U87MG, T98G, and A172MG were grown in Dulbecco’s Modified Eagle Medium (DMEM, Gibco) whereas ADF was cultured in RPMI 1640 (Corning) supplemented with 10% FBS (Lonza), 100U/mL penicillin and 100mg/mL streptomycin (Life Technologies) at 37°C and 5% CO2. GBM cell lines U87MG, T98G, and A172MG were grown in Dulbecco’s Modified Eagle Medium (DMEM, Gibco) whereas ADF was cultured in RPMI 1640 (Corning) supplemented with 10% FBS (Lonza), 100U/mL penicillin and 100mg/mL streptomycin (Life Technologies) at 37°C and 5% CO_2_

### Molecular modulation of TSPO

The downregulation of TSPO in the U87MG cell line (U87MG-TSPO) was achieved using the pGIPZ shRNA vector: clone ID V3LH_331646, target sequence: 5′-TGAGTGTGGTCGTGAAGGC-3, purchased from Open Biosystems (Huntsville, AL, USA). The plasmid contains the TurboGFP (tGFP) reporter for visual tracking of transduction and expression. Transfection was performed using standard Ca^2+^ phosphate-based protocol as previously demonstrated (Campanella et al., 2004; Morelli et al., 2003). After transfection, cells were maintained in media supplemented with 3μg/ml puromycin (SERVA Electrophoresis GmbH, 33835) for selection of the GFP-positive cells. TSPO downregulation was further confirmed by immunoblot analysis.

The overexpression of TSPO in the U87MG cell line (U87MG+TSPO) was performed using the TSPO encoding plasmid from (Origene RG220107) which contains the C-terminal MYC/DDK tag, reporter for visual expression. Transfection was performed using Lipofectamine 2000 as per the manufacturer’s instructions. After transfection, selection was made using G418 (700 μg/μL) and Western Blot analysis enrolled to confirm TSPO overexpression.

### Treatments

The compounds Temozolomide (Sigma, T2577), PK11195 (Enzo Life Technologies, BML-CM 118), Etifoxine (Sigma, SML 0272), Lovastatin (Enzo, BML-G226), Cerivastatin (Sigma, SML0005) and MF-438 (Sigma, 569406) have been enrolled in the analysis at the concentration and time indicated in the specific experiments.

### Cell viability analyses

Cell viability was measured using crystal violet solution (Sigma, HT90132) or red fluor PI (propidium Iodide) as previously described (Feoktistova et al., 2016). Mitochondrial metabolic activity, as the function of cell health was detected via 3-(4,5-dimethylthiazol-2-yl)-2,5-diphenyltetrazolium bromide (MTT, Invitrogen, M6494) assay (Ricci et al., 2013). Cell Titer Glo 3D (Promega, G9683) was used to evaluate the cellular viability of ADF in 3D cultures as previously demonstrated (Ricci et al., 2013).

### Sphere formation assay

The U87MG cells were seeded at 2000 cells/well in the PrimeSurface® culture ware ultra-low attachment (ULA) U bottom microplates and allowed to grow for 4 days. On day 4, the spheroids underwent treatments with the conditions of interest.

Sphere propagation assays were performed as previously described (Noto et al., 2017). Briefly, single-cell preparation (1000 cells/well) of ADF cell lines were suspended in an appropriate amount of sphere-forming medium: serum-free DMEM/F12 (Gibco) supplemented with basic Fibroblast Growth Factor (bFGF, Sigma Aldrich, F0291), Epidermal Growth Factor (EGF, Sigma Aldrich, E4127), insulin (Sigma Aldrich, 9011-M), glucose (Sigma Aldrich, G7021), heparin (Sigma, 2106), B27 (Gibco, 17504001) and plated into a 96-well ultra-low adherent plate (Costar) to form spheres. On day 4, the sphere-forming efficiency was evaluated, and the experimental conditions of interest were applied.

### Subcellular fractioning and Western blot analysis

GBM cells were washed in Phosphate Buffer Saline (PBS, Gibco), collected by gentle scraping, and incubated in ice-cold RIPA buffer (50mM Tris–HCl, 150mM NaCl, 1% (v/v) TritonX100, pH 8.0) containing protease and the phosphatase inhibitor for 30 min on ice. Total lysates were centrifuged at 13,000g at 4◦C for 20 min. The supernatant was collected to detect protein concentration. Subcellular fractionation was performed with the Rapid Efficient And Practical (REAP) method(Suzuki et al., 2010). GBM cells were incubated in Reap Lysis Buffer (PBS 1X and 0,1% Nonidet P-40 Substitute) for 3 min on ice. Lysates were centrifuged at 2200g for 5 min at 4°C. The pellet corresponding to the nuclear fraction was washed, centrifuged 3 times at 2200rpm for 5 min at 4°C and sonicated 3 times for 5 sec, amplitude 80. Protein concentration was estimated using the Bradford reagent (Biorad, 5000006), and 20-40μg of total proteins was mixed with Laemmli sample buffer (Biorad) and boiled at 95°C for 2 min. Proteins were resolved by Sodium Dodecyl Sulfate Polyacrylamide gel (SDS–PAGE) and transferred to PVDF (Merk Millipore) membranes. Membranes were blocked in 5% Bovine Serum Albumine (BSA) (Sigma, A7030) or 5% non-fat dry milk (Applichem, A0830) in 1×Tris buffer saline (TBS) (25 mMTris, 0.15 M NaCl) containing 0.05% Tween-20 (Sigma, P9416) (TBST, pH 7.5) for 1h, probed with appropriately diluted primary (GAPDH (Abcam, ab9485), MTCO1 (Abcam, ab203912), TSPO (Abcam, ab118913), OPA1 (BD, 612607), MFN1 (Santa Cruz. Sc-166644), HSP90 (Santa Cruz, sc-69703) Lamin A/C (Cell signalling, 4777), Lamin B2 (Thermofisher Scientific, 33-2100), ATPβ (Abcam, ab14730), Actin (Sigma-Aldrich, A2066), MnSOD (Millipore, 06-984), SREBP-1 (Santa Cruz, sc-13551), YAP (Santa Cruz, sc-101199) and LXR (Abcam, ab3585)) overnight at 4°C washed in TBST (3×10 min at RT) and then probed with the corresponding peroxidase-conjugated secondary antibody (Biorad) for 1h at RT. After further washing in TBST (3×10min at RT), proteins were detected using an ECL detection kit (Merk Millipore) via Amersham Imager 600. Immunoreactive bands were analysed by performing densitometry analysis with ImageJ software (NIH, Bethesda, MD, United States).

### RNA extraction, reverse transcription, and quantitative real-time PCR

U87MG cells were lysed using GENEAID - Total RNA Mini Kit (Geneaid Biotech Ltd, RBD100), and total RNA was extracted using TAKARA - PrimeScript RT Reagent Kit (Takara, RR037A) according to the manufacturer’s instructions. In all, 1μg of RNA was retro-transcribed using SensiFAST™ cDNA Synthesis Kit (Bioline) and used in quantitative RT-PCR (qPCR) experiment, using SensiFAST™ SYBR Hi-ROX Kit (Bioline) following the manufacturer’s instructions. Thermocycling consisted of an initial polymerase activation step at 98 °C for 5 min, and amplification was performed with 35 cycles of 95 °C for 15 s, 68 °C for 10 s, and 72 °C for 20 s with data acquisition at this stage and the reaction finished by the built-in melt curve. The relative amounts of mRNA were calculated by using the comparative Ct method. Primers for GAPDH: 5’-TCATGGGTGTGAACCATGAGAA-3’ (forward) and 5’-GGCATGGACTGTGGTCATGAG-3’ (reverse); TSPO: 5’-TCTTTGGTGCCCGACAAATGG-3’ (forward) and 5’-TCAACTACTGCGTATGGCGGGACAACC-3’ (reverse); CYP11: 5’-CCGTGACCCTGCAGAGATAT-3’ (forward) and 5’-TGGTCATCTCTAGCTCAGCG-3’ (reverse). Gene expression was normalized to GAPDH mRNA content.

The total RNA of ADF was extracted and isolated by TRIzol (Thermo Fisher, 15596018) following the manufacturer’s instructions (D’Amico et al., 2013). Quantitative RT-PCR was performed as previously described (Noto et al., 2017; Pisanu et al., 2014). Gene expression was normalised to H3 mRNA content.

### Immunofluorescences

#### On tissue

Tissue microarrays including formalin-fixed and paraffin-embedded cores of healthy brain and WHO grade 1 to 4 human astrocytoma were purchased from US Biomax (US Biomax, Inc.Rockville, MD 20849, United States) (codes GL208 and GL722). Following PBS washing, primary human GBM sections were processed for antigen retrieval and incubated at 80°C for 30m in citrate buffer (10mM Sodium Citrate dihydrate (Sigma, W302600), 0.05% Tween-20 (Sigma, P9416), pH 6.0). Washes were flowingly carried out in PBS-T (0.05% Tween-20 in PBS). Permeabilization was carried out for 15 min at 4°C in 20mM HEPES, 300mM Sucrose (Sigma, S0389), 50mM NaCl, 0.05% Triton-X100 (Sigma, T8787), 3mM MgCl2, 0.05% Sodium Azide (Sigma, S2002). Blocking – 1h at room temperature in 10% horse serum (Thermo Fisher), 1% w/v Bovine Serum Albumin (BSA, Applichem, A1391) in PBS. TSPO (Abcam, ab118913) and NFkβ (Cell Signaling, #6956) primary antibody incubation were carried out overnight at 4°C followed by the secondary antibodies anti-rabbit (Invitrogen, A-11012) and anti-mouse (Invitrogen, A-11032) in a blocking solution for an hour at RT, while shaking. Slices were washed 4-5x in PBS-T and 3x in PBS, before a 5 min incubation with 600nM DAPI solution (Applichem, A4099) for nuclei visualization. Excess DAPI was cleared with 3×5 min washes in PBS and slices were then mounted on glass slides (VWR) using a ProLong Gold antifade reagent (Invitrogen, P36930), covered with 22 mm glass coverslips. The slices were then imaged using an Olympus Fluoview 1000 Confocal Laser Scanning Microscope using objective 60x oil (N.A. 1,35) and laser 405nm for DAPI fluorescence, 488nm for green fluorescence and 543nm for red fluorescence. All the images were processed with Imaris software (Bitplane, Switzerland) for 3d rendering and isosurfaces and analysed with ImageJ software for colocalization analysis. All images were analysed by ImageJ software (NIH, Bethesda, MD, United States).

#### On cells

Following treatment GBM cells were washed once with Phosphate Buffered Saline (PBS) and fixed with 4% paraformaldehyde (PFA) following an incubation of 10 minutes. Cells were subsequently washed 3 times for 5 minutes each with PBS and permeabilized with a 0.1% Triton-X-100 (Applichem, A4975)/PBS solution for 10-20 minutes. After another round of washes cells were blocked for 1 hour in a solution of PBS containing 3% w/v BSA. Primary antibodies ATPβ (Abcam, ab14730) and TSPO (Cell Signalling, 70358), were then incubated overnight at 4°C in a blocking solution. After 3 washes, the secondary antibody was incubated for 1 hour in a blocking solution: anti-mouse (Invitrogen, A-11094) and anti-rabbit (Invitrogen, A-11012). After 1 hour at room temperature away from light, the coverslips were washed and mounted onto glass slides using a 4’,6’-diamidino-2-phenylindole (DAPI)-containing mounting medium (ab104139). The coverslips were then imaged on an Olympus Fluoview 1000 Confocal Laser Scanning Microscope using objective 60x oil (N.A. 1,35) and laser 405nm for DAPI fluorescence, 488nm for green fluorescence and 543nm for red fluorescence. All the images were processed with Imaris software (Biplane, Switzerland) for 3d rendering and isosurfaces and analysed with ImageJ software.

For the detection of cellular cholesterol, we used a cell-based cholesterol assay kit (ab133116; Abcam, Cambridge, MA). Briefly, U87MG cells were fixed with PFA (4%) and stained with filipin III following the manufacturer’s instructions. Cells were visualized using a microscope Zeiss Axio observer 7. All images were analysed and processed by ImageJ software (NIH, Bethesda, MD, United States). Mitochondrial fragmentation was detected using Mitochondrial Network Analysis (MiNA) ImageJ plug-ins as previously described (Valente et al., 2017).

### Immunoprecipitation

The nuclear fractionation of U87MG cells was obtained as previously described (Suzuki et al., 2010). Protein concentration was estimated using the Bradford reagent (Biorad, 5000006) so that 1.5 mg of nuclear extracts were incubated with anti-TSPO (Cell Signalling, #70358) and rotated at 4 °C overnight. The day after, the Dynabeads™ M-280 Sheep Anti-Rabbit IgG (11203D) were equilibrated in Reap Buffer (0.1% NP40 in PBS) and collected by magnet 3 times. Then 20-30µl of beads were added to each sample and rotated at 4 °C for 2h. After that the suspension was put in the magnet: the supernatant was harvested as unbound whereas the rest bead slurry was washed wash 3 times with Reap Buffer. After the last wash, the beads were completely dried and resuspended with 2x Laemmli Buffer (Biorad) and boiled at 95°C for 5 min. As a negative control, the beads were incubated with the anti-TSPO antibody and rotated at 4°C for 2h.

### Transmission Electron microscopy (TEM)

U87MG cells (DMSO and treated) were fixed at RT with 2.5% glutaraldehyde (v/v) in 0.1 M sodium cacodylate buffer (pH 7.4) for 4 hours at 4°C. The cells were pelleted by centrifugation, washed in buffer, and post-fixed for 1 h with 1 % osmium tetroxide in 0.1 M sodium cacodylate buffer. Samples were thoroughly rinsed, dehydrated in a graded series of ethanol, and embedded in epoxy resin (TAAB 812). Ultrathin sections (60-70 nm) were cut using a Leica UC7 ultramicrotome mounted on 200 mesh copper grids and contrasted using Uranyless (TAAB). Samples were observed under a transmission electron microscope JEOL JEM 2100 Plus (Japan Electron Optics Laboratory Co. Ltd. Tokyo, Japan). Images were captured digitally with a digital camera TVIPS (Tietz Video and Image Processing Systems GmbH. Gauting, Germany). The mito/nuclear interactions were detected as previously demonstrated (Desai et al., 2020; (Lam et al., 2021).

### Analysis of primary and recurrent human glioblastoma

We investigated the primary and recurrent glioblastoma of 5 patients who progressed during chemotherapy in a cohort of 85 who underwent maximally safe tumour removal and adjuvant radiotherapy and TMZ treatment between January 2015 and December 2016. At the time of this study, all patients passed away from tumour exacerbation. The use of retrospective, archived, anonymised tissue samples of deceased patients has been duly approved by the Human Research Authority (IRAS 244538) and the Research Ethic Committee assessing the project (19/NE/186). The expression of TSPO was assessed in the primary tumour and recurrence from adult patients with supratentorial, hemispheric glioblastoma who progressed after radiotherapy and during the cycles of TMZ. Both primary tumour and recurrence were operated on by one neurosurgeon (PdU) and underwent maximally safe removal with 5-ALA guided intra-operative microscopy. The original slides were reviewed by a neuropathologist (FR) to identify the most representative areas for immunohistochemistry and the histotype redefined according to the criteria of the 2021 World Health Organisation Classification of Tumours of the Central Nervous System. The tissue was fixed in formalin and routinely processed to paraffin embedding for diagnostic purposes. All cases were routinely tested for IDH1 (R132H) using immunohistochemistry (anti-IDH1 (R132 Dianova, Hamburg, Germany monoclonal H09, 1:200) and non-canonical IDH1 mutations and IDH2 mutations using next-generation sequencing (sequence analysis was carried out following PCR enrichment using a QIAseq Targeted DNA custom panel, Germantown, MD, USA and Illumina Next Generation Sequencing, Illumina, San Diego CA, USA). MGMT gene promoter methylation was tested using pyrosequencing (check details). MGMT resulted in unmethylated in the five selected primary tumours. The immunoreaction for TSPO and YAP1 was performed on a BenchMark ULTRA Slide Staining System (Roche Diagnostics, Indianapolis, IN, USA) using the anti-TSPO, goat polyclonal (Abnova, Walnut, CA USA) and the mouse monoclonal anti-YAP1 (Santa Cruz Biotechnology).

### Bioinformatics analysis

To investigate the impact of TSPO expression on progression-free survival (PFS), two RNA datasets were retrieved from the UCSC Xena system and analysed: the TCGA GBM and LGG cohorts. Normalized RNA expression values and survival clinical data were downloaded from the UCSC Xena system for TCGA glioblastoma (GBM; RNA-seq n=160)11 and low-grade glioma (LGG n=514; RNA-seq)12 cohorts. Each of the two cohorts was split on three groups of equal13 size according to the quantiles of gene expression level (cut2(), R “Hmisc” package). Then, for each group, we fitted a survival model according to the progression-free survival (PFS) time (R “survival” package).

To assess the level of Hippo-YAP pathway activation using RNA data for the TCGA samples (GBM and LGG cohorts), we used a relative level of TEAD2 transcription factor activation measured by the expression of its downstream target genes with the Dorothea pipeline (Garcia-Alonso et al., 2019). Only regulons with the A-D level of confidence were used. The relative activity of TEAD2 was assessed using Viper algorithm15 with the following settings: method = "scale", minsize = 10, nes=T, pleiotropy=T. Finally, the normalized enrichment score of inferred TEAD2 activity was correlated with the gene expression of the TSPO gene.

### MGMT Promoter Methylation assay

MagPurix extraction kit ZP02005 (Zinexts, Life Science Corporation, New Taipei City, Taiwan) with Zinexts automated system was used for DNA extraction from cell cultures. DNA quantification was performed with the Qubit® dsDNA BR Assay Kit (Thermo Fisher Scientific, Waltham, USA). A maximum of 200 ng of DNA per sample underwent Bisulfite conversion using the Zymo EZ Methylation Kit (Zymo Research Irvine, USA).

We used the MGMT Promoter Methylation Detection Kit (EntroGen, CA, USA) following the manufacturer’s instructions. In detail, the kit covers CpG sites between 75 and 86 (12 sites) starting at chr10:129,467,243 (hg38 genome build), to calculate MGMT promoter methylation status within the DMR2 region, located in exon 1, just after the transcription start site of the MGMT gene. To distinguish between methylated and non-methylated cytosines in the MGMT gene, the assay uses specific primers to amplify only methylated MGMT CpG sites and amplifies an endogenous control gene (Actin B) about which the extent of MGMT promoter methylation is calculated, using a semi-quantitative real-time PCR QuantStudio 7 Pro system (Applied Biosystems™ Waltham, USA).

### Statistical analysis

Statistical analyses were performed using GraphPad Prism 9 software, choosing the most appropriate test. The number of independent experiments and the number of replicates used to perform statistics is ≥ three. Unpaired Student’s t-test and ordinary one or two-way ANOVA followed by the Bonferroni Test were used for cellular correlation and multiple comparisons. Post-hoc test was conducted only if F achieved P<0.05. Data are presented as means ± standard error of the mean (SEM). Statistical significance was declared at p ≤ 0.05. The asterisks describe different value levels of statistical significance: ****P<0.0001, ***P<0.001, **P< 0.01 and *P<0.05.

## Results

### TMZ reprograms mitochondria and counteracts mitophagy in glioblastoma cells

U87MG cells were exposed to 100μM TMZ for 3h followed by a recovery time of 3 (D3) (**Fig.1A**), 5 (D5) and 7 (D7) days as previously published (Filippi-Chiela et al., 2015). Cumulative population doubling was lower in the TMZ-treated cells as reported in **Suppl.Fig.1A**. Mitochondrial membrane potential (ΔΨ_m_) and mitochondrial mass were evaluated on D3, D5 and D7 using the fluorescent probes JC-1 (**Suppl. Fig.1B**) reporting an increase matched by one on the Reactive Oxygen Species (ROS) at D5 (**Suppl.Fig.1C**). To evaluate the mitochondrial respiration, the rate of mitochondrial oxygen consumption was tested using high-resolution respirometry (HRR) Oxygraph-2k which recorded a significant increase on D5 (**Suppl. Fig.1D**). Furthermore, by assessing the flux control ratio (FCR), we observed that the TMZ pre-treated cells have a lower leak and higher spare reserve capacity in their mitochondria compared to untreated cells (**Suppl. Fig.1E**) thus implying a metabolic switch towards a more oxidative state. This was also confirmed by the U87MG cells being more sensitive to OXPHOS inhibitor KCN (137μM and 40μM respectively to control and TMZ-treated cells) than glycolysis inhibitor 2DG (4mM) (**Suppl. Fig.1F**). To inspect autophagy triggered by TMZ as previously demonstrated (Filippi-Chiela et al., 2015), we enrolled the fluorescent dye Acridine Orange (AO)(Murugan & Amaravadi, 2016) to map the late stages of autophagy finding a tangible increase of signal at D3 and D5 (**Suppl. Fig.1G-H**). This prompted us to monitor autophagy throughout the prolonged exposure to the therapeutic agent (Le Calvé et al., 2010) namely after 72 and 120h of treatment (**Suppl. Fig.1I-J**). In these conditions, following inhibition of the Autophagy flux via NH_4_Cl, we monitored the LC3 level via immunoblotting analysis recording a rise in autophagy via TMZ (Filippi-Chiela et al., 2015) (**Suppl. Fig.1I-J**) corroborated by imaging means(Itakura & Mizushima, 2010) (**Suppl. Fig.2A-B**).

**Figure 1.**
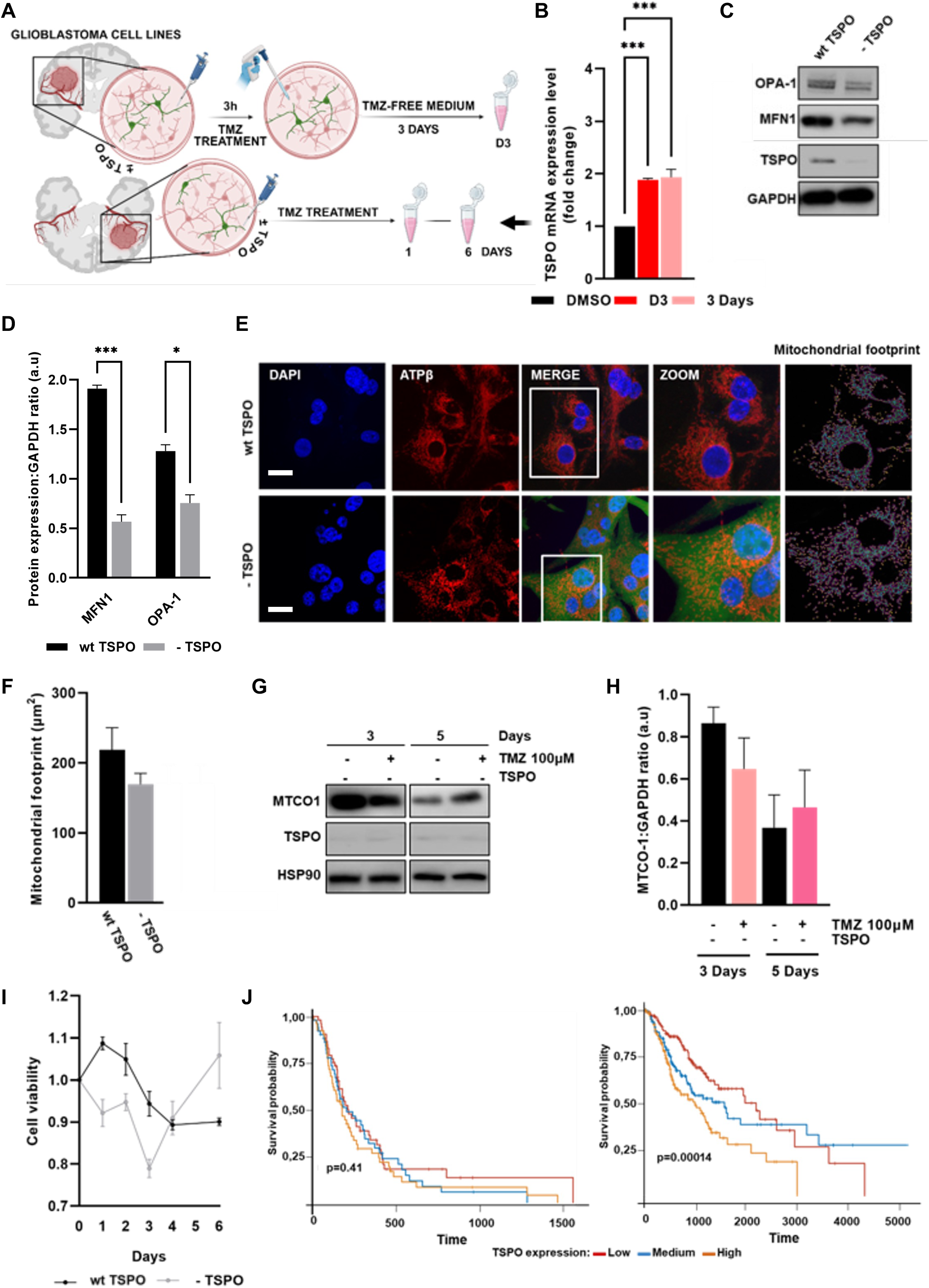
Fragmentation of the mitochondrial network is required for TMZ-induced demise. **(A)** Schematic of the experimental design for u87MG cells treatment with TMZ. **(B)** q-RNA analysis of TSPO in D3 and 72 hours protocols of TMZ treatment. **(C)** Immunoblotting of OPA-1, MFN-, MTCO-1 and TSPO in lysates of u87MG + and - TSPO normalized to HSP90 and quantified in **(D)**. **(E)** Confocal images of the immunocytochemical analysis of the mitochondrial ATPβ (red) in U87MG + TSPO and -TSPO, U87MG – TSPO (bar=100μ). **(F)** reports their mitochondrial network footprint analysis via MiNA plug-in. **(G)** Immunoblotting of MTCO-1 and TSPO in U87MG – TSPO TMZ-treated for 72 (3 days) and 120hours (5 days) (100μM) with densitometry analysis-normalized over HSP90- in **(H)**. **(I)** Cell viability was evaluated via crystal violet, in both U87MG + and – TSPO at different time points of TMZ treatment (100μM). **(J)** Kaplan–Meier plots for progression-free survival for TCGA patients with glioblastoma (GBM, n=160) and low-grade glioma (LGG, n=514) stratified by the level of *TSPO* gene expression (two groups of expression level for upper panels and three groups of expression level for bottom panels). P-values reflect the outcomes of log-rank tests for the survival model. All data are represented as mean±sem. *p≤0.05; **p≤0.01; ***p≤0.001

**Figure 2.**
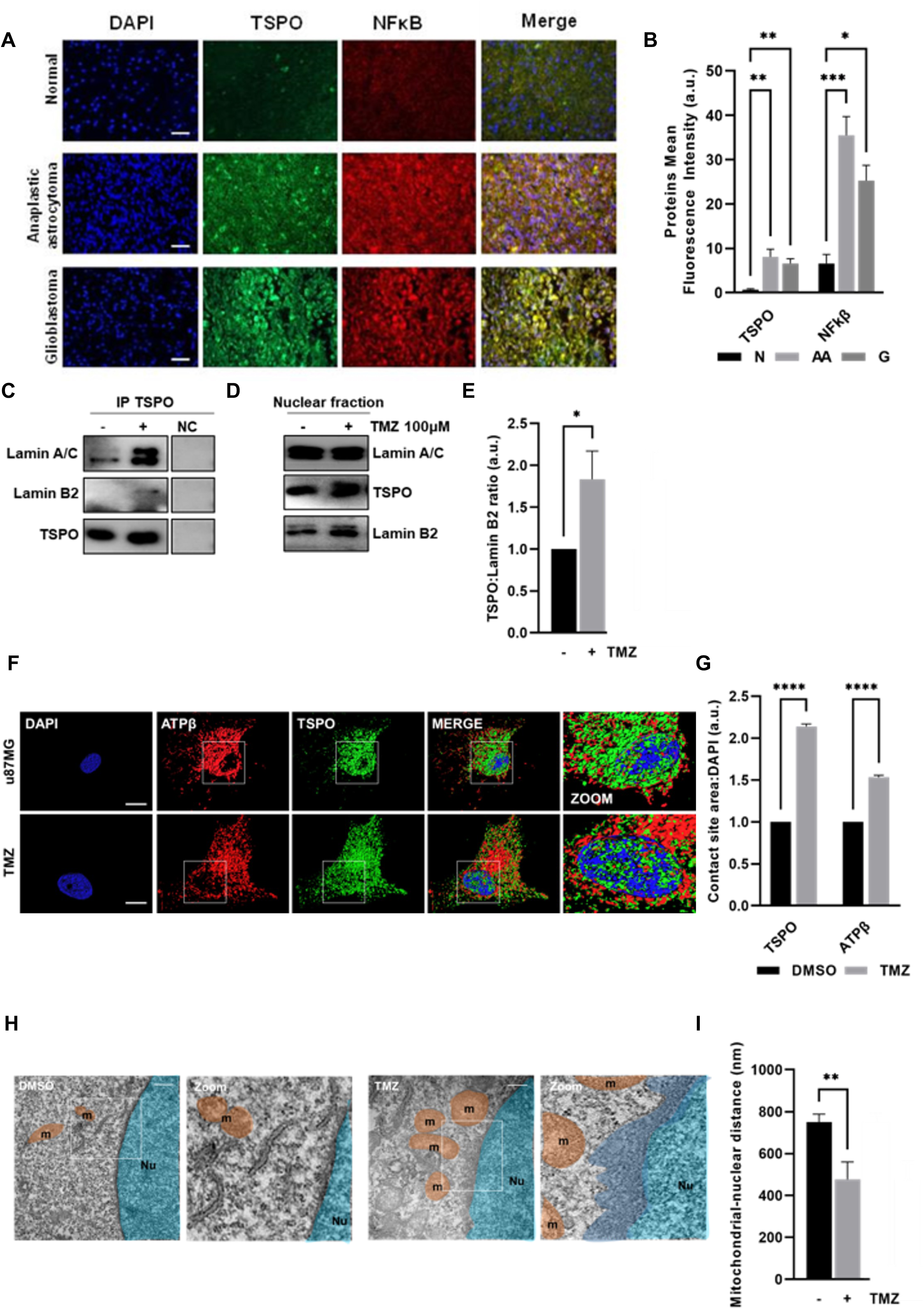
TMZ primes mitochondrial redistribution on the nucleus. **(A)** Immunohistochemical analyses of patients (N=Normal) derived diffuse astrocytoma (AA= anaplastic astrocytoma) and glioblastoma (G) sections stained with TSPO (green) and NF-κB (red) (bar=50µm) quantified in **(B)**. **(C)** Co-Immunoprecipitation of TSPO and Lamin A/C or B2 in TMZ-treated U87MG at 24h-timepoint (TMZ, 100 μM). **(D)** Immunoblotting (IP input) of TSPO, Lamin A/C and B_2_, in nuclear fractions of U87MG TMZ-treated (100μM) for 24h with densitometry analysis-normalized over Lamin B2: quantified in **(E).** (**F**) Confocal microscopy and isosurface analysis of ATPβ and TSPO in TMZ-treated (100μM). The mitochondria-nucleus contact site is represented in **(G)**. **(H)** TEM micrographs of TMZ-treated U87MG (24h-timepoint, 100μM) and relative control highlighting the physical interaction between mitochondria (orange) and the nucleus (blue) quantified in (**I**) (bar=5µm). All data are represented as mean±sem. *p≤0.05; **p≤0.01; ***p≤0.001; ****p≤0.0001

Then, we tested the levels of TSPO, a stress-driven inhibitor of mitophagy (Gatliff et al., 2014) (i) and the mitochondrially encoded cytochrome c oxidase I (MTCO-1) protein (ii) which works as a bona fide read-out of mitochondrial processing via autophagy. In the total lysates of U87MG cells attained after either acute or chronic treatment with TMZ (**Fig.1A**), we observed an-accumulation of both TSPO and MTCO-1 protein levels (**Suppl. Fig.1K-L**). This was indicative of a blockage in the mitophagy flux which was further assessed by performing confocal microscopy analysis of the co-localization between either TSPO (green) and LC3 (red) (**Suppl. Fig.2A-B**) or (ii) the mitochondrial ATP synthase β-subunit (ATPβ, green) and lysosomes via the Lysotracker Red (**Suppl. Fig.2C-D**). Mitophagy is therefore repressed by TMZ for the concomitant stabilization of TSPO whose level rapidly increases when U87MG cells are challenged with the chemotherapeutic (**Fig. 1B**).

### Stabilization of TSPO is paired with mitophagy repression and formation of NAM

To inform the role of TSPO in dictating susceptibility or resistance to TMZ we devised a line stably downregulated for the mitochondrial protein: U87MG-TSPO (**Fig. 1C**). The U87MG-TSPO lines showed a high level of mitochondrial fragmentation (**Fig. 1E, F**) as confirmed by the processing of pro-fusion proteins MFN1, OPA-1 (**Fig.1C, D**). Pruning of the mitochondrial network following downregulation of TSPO, promoted by mitophagy activation, well read by the loss of the mitochondrial MTCO-1 (**Fig.1G-H**), linked to a modification of TMZ mediated cytotoxicity (**Fig.1I)**. To frame the clinical relevance of these results, we investigated the impact of TSPO expression on the survival probability of patients diagnosed with Glioblastoma (GBM) or Low-Grade Glioma (LGG). Out of these two populations, mining from specific databases indicated in the methods, we assessed 160 samples and 514 samples of RNASeq mined from databases from these two populations (**Fig.1J**). The analysis revealed that the differential expression level of TSPO does not have a significant impact on the prognosis of GBM. In contrast, a statistically significant association was found between reduced TSPO level and good prognosis in LGG.

The over-expression of TSPO in high-grade gliomas is well documented in the literature (Ammer et al., 2020; Roncaroli et al., 2016) and we corroborated this further by assessing tissue sections of a normal brain (i), anaplastic astrocytoma (WHO grade 3) (ii) and glioblastoma (WHO grade 4) (iii). This experiment documented in **Fig.2A-B** not only confirmed the positive correlation between TSPO and the aggressive tumour grades but also revealed another one between TSPO and NF-kB. This transcription factor not only indicates an active exploitation of the mitochondrial retrograde response (Desai et al., 2020) but also the formation of contacts between the two organelles (Singh et al., 2021).

Using co-immunoprecipitation (Co-IP), immunohistochemistry, and ultrastructural imaging validations we assessed the perinuclear clustering of mitochondria following TMZ treatment. These analyses unveiled an interaction between TSPO and two nuclear Lamins (A/C and B2) in U87MG cells following 24h treatment with TMZ (100μM) (**Fig.2C, D**) along with an increase of TSPO in the nuclear fraction (**Fig2.E**). This data was further corroborated via immunostaining (**Fig.2F, G**) and transmission electron microscopy (TEM) (**Fig.2H, I**).

The redistribution of mitochondria towards the nucleus (**Fig.2D-I**), facilitated by the repression of mitophagy (**Suppl. Fig2C, D**), and the consequent re-adaptation of the nuclear membrane towards the mitochondria would scale MRR hence resistance to TMZ.

TMZ-mediated cytotoxicity would therefore preserved if not potentiated if TSPO expression was downregulated or its function impaired via the protein ligands. Furthermore, we also speculated that an equal outcome could be achieved by lowering the level of cholesterol which feeds the TSPO pathway. All this was tested in the next set of experiments.

### Pharmacological modulation of TSPO potentiates TMZ-mediated cell death

We assessed the viability (Maaser et al., 2001) of U87MG cells following modulation of their TSPO level by overexpressing the protein (+TSPO) or ablating its expression (-TSPO). The survival rate was also tested in response to TMZ treatment alone (100µM) or in combination with either the cholesterol-lowering drug Lovastatin (10µM) or the TSPO ligand Etifoxine (30µM) (**Fig.3B**). Subsequently, we extended the analysis to A172MG, T98G and ADF cell lines (Bonfigli et al., 2017; Giorgini et al., 2007) to validate the findings in models which bear alternative levels of O6-methylguanine (O6-MeG)-DNA methyltransferase (MGMT) expression (Yachi et al., 2008) which we confirmed in **Suppl. Fig.2G**. MGMT protects cells from the lethal effects of chemotherapy with DNA alkylating activity. In A172MG cells, which are devoid of the MGMT-like the U87MG ones- we tested the susceptibility to TMZ-induced cell death following co-treatment with Etifoxine and Lovastatin (**Fig.3B**).

**Figure 3.**
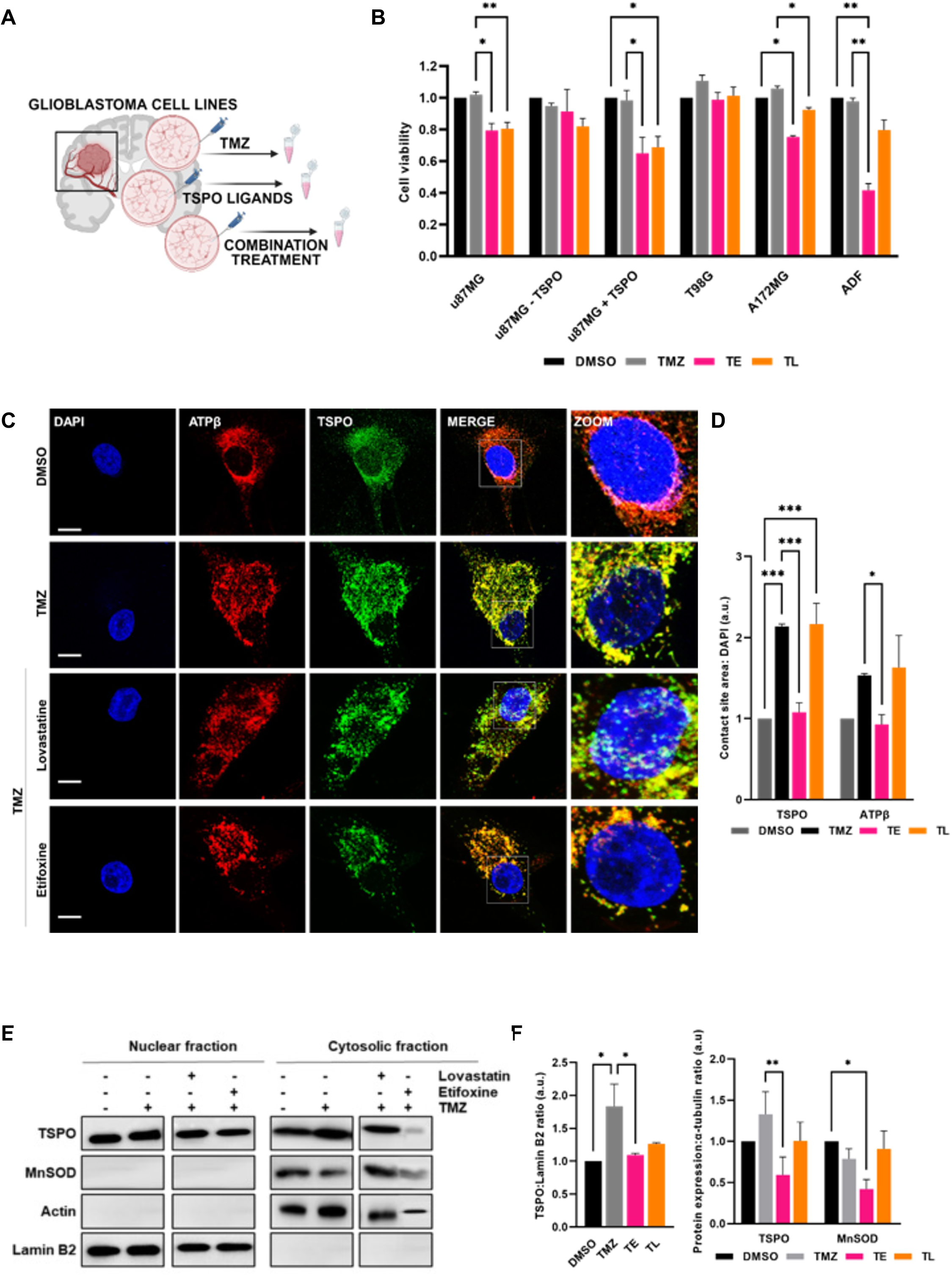
Pharmacological regulation of TSPO and cholesterol synthesis co-adjuvant TMZ induced cell death. **(A)** Schematic of the experimental design for glioblastoma cells treatment with TMZ and TSPO ligands. **(B)** Cell viability was evaluated via crystal violet in u87MG, u87MG + TSPO, u87MG - TSPO, T98G, A172MG and ADF treated with Etifoxine (30μM) and Lovastatin (10μM) in combination with TMZ (100μM) for 24 hours. **(C)** Isosurfaces derived from confocal images of immunocytochemical analyses of TSPO (green) and ATPβ (red) at 24h-time point of TMZ treatment (100μM) combined with Lovastatin (10μM) and Etifoxine (30μM) (bar=100μ). **(D)** Quantification of the mito-nuclear contacts. **(E)** Immunoblotting of TSPO, MnSOD, Actin and Lamin B2 in nuclear and cytosolic fractions of TMZ-treated U87MG (24h-timepoint, 100μM) alone and combined with Lovastatin(10μM) or Etifoxine (30μM). **(F)** Reports densitometry analysis of TSPO and MnSOD levels. All data are represented as mean±sem. *p≤0.05; **p≤0.01; ***p≤0.001

The TSPO ligand PK11195 (Levin et al., 2005) was also tested (**Suppl. Fig.3A, B**) but we focused on Etifoxine (**Suppl. Fig.4C**) which proved to be capable of generating a greater potentiating effect on TMZ-mediated cell death (**Suppl. Fig.4E**).

**Figure 4.**
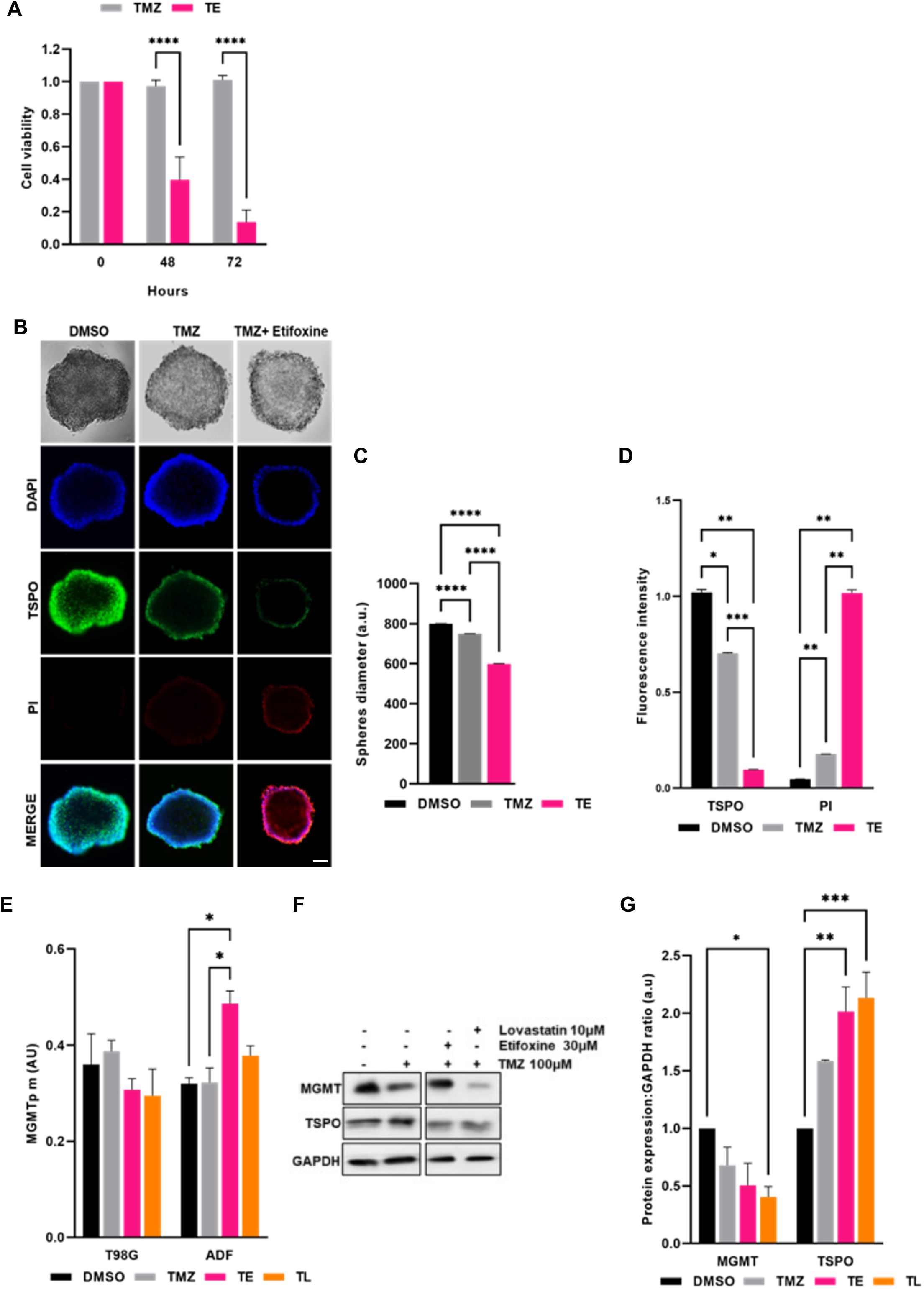
TSPO determines cell death and MGMT methylation in glioblastoma cells. **(A)** Cell viability was evaluated via crystal violet in U87MG cells treated with Etifoxine (30μM) and in combination with TMZ (100μM) for 48 and 72 hours. **(B)** Representative images of U87MG spheroids stained with Dapi, PI and antibody for TSPO at resting conditions, following TMZ alone and combined with Etifoxine (TMZ+Etifoxine). Histograms reporting quantification of spheroids diameter in the conditions of analysis (**C**) and relative changes in fluorescent intensity (**D**). **(E)** Percentage of MGMT promoter methylation (MGMTp meth%) in t98G and ADF cell lines. **(F)** Immunoblotting of MGMT and TSPO in ADF total lysates treated with TMZ (100μM) alone or combined with Lovastatin (10μM) and Etifoxine (30μM) for 24 hours. The graph in panel (**G**) shows the relative densitometry of MGMT and TSPO normalized to GAPDH. All data are represented as mean±sem. *p≤0.05; **p≤0.01; ***p≤0.001

In these experimental conditions, we also checked the redistribution of mitochondria by immunostaining the subunit β of the F1-Fo-ATPsynthase and TSPO after TMZ treatment alone and in combination with Etifoxine or Lovastatin (**Fig.3C, D**). The analysis was furthered by using nuclear and cytosolic fractions of the U87MG cells and monitoring TSPO expression level via immunoblotting analysis following the specific treatments (**Fig.3 E, F**). This confirmed a neat nuclear accumulation of TSPO following TMZ-mediated chemical stress likewise a reduction of its content in the cytosolic portion when combined with Etifoxine or Lovastatin. The combined treatment between TMZ and Etifoxine tangibly impacts cellular growth at 48 and 72h of co-treatment (**Fig 4A**).

Aware of the limitations of the 2D way of cellular growth in securing a pharmacological inferring we tested the cell modulation attained with the TSPO ligand Etifoxine in a 3D analysis. Spheroids of U87MG were therefore attained and assessed for dimension and percentage of dead cells following treatment with TMZ (100μM) or a combination of the chemotherapeutic agent with Etifoxine (30μM) for 72 hours (**Fig. 4B-D**).

Out of the MGMT-positive cell lines T98G and ADF, only the latter showed susceptibility to the treatment (**Fig.3B**). Accordingly, in both cells we evaluated the methylation status of the *MGMT* promoter (as a signature of epigenetic silencing) (Yachi et al., 2008) (**Fig.4E**) and the actual expression level of the enzyme following treatment with Etifoxine and the cholesterol-lowering drug Lovastatin (**Fig. 4F, G**). Both conditions increased methylation which was met by a reduction of MGMT expression otherwise enhanced by TMZ treatment (**Fig.4F-G)**. However, of the two approaches the prominent effect was visible with the combination of Lovastatin and TMZ.

The upstream effects of both these co-treatments have been further assessed starting from the one on cholesterol signalling and metabolism. Considering that Lovastatin impairs cholesterol biosynthesis (Krukemyer & Talbert, 1987) and Etifoxine promotes cholesterol influx in the mitochondria via TSPO (Strobbe & Campanella, 2018), we hypothesised that transport and metabolism of the lipid might be affected hence its redistribution taking part in the resistance mechanisms exploited by the glioblastoma cells exposed to TMZ.

### TMZ combined with Etifoxine redesigns the nuclear localization of Transcription Factors

Initially, we checked the intracellular localization of cholesterol via the fluorescent compound filipin in U87MG cells (**Fig.5A**). This showed a redistribution of the lipid, from the nuclear region of the cell to the plasma membrane when TMZ (100μM) was combined with Etifoxine (30μM, 24h) and Lovastatin (10μM, 24h). Subsequently, we detected the mRNA level of the mitochondrial cytochrome P450 side chain cleavage (CYP11A1) enzyme involved in the conversion of cholesterol to pregnenolone after treatments (**Fig.5B**). This was greater when TMZ was combined with Etifoxine thus implying a sustained rate of cholesterol conversion leading to a reduction of the lipid intracellularly hence on is mediated pathogenicity in the very cell type (Sorrentino et al., 2014).

**Figure 5.**
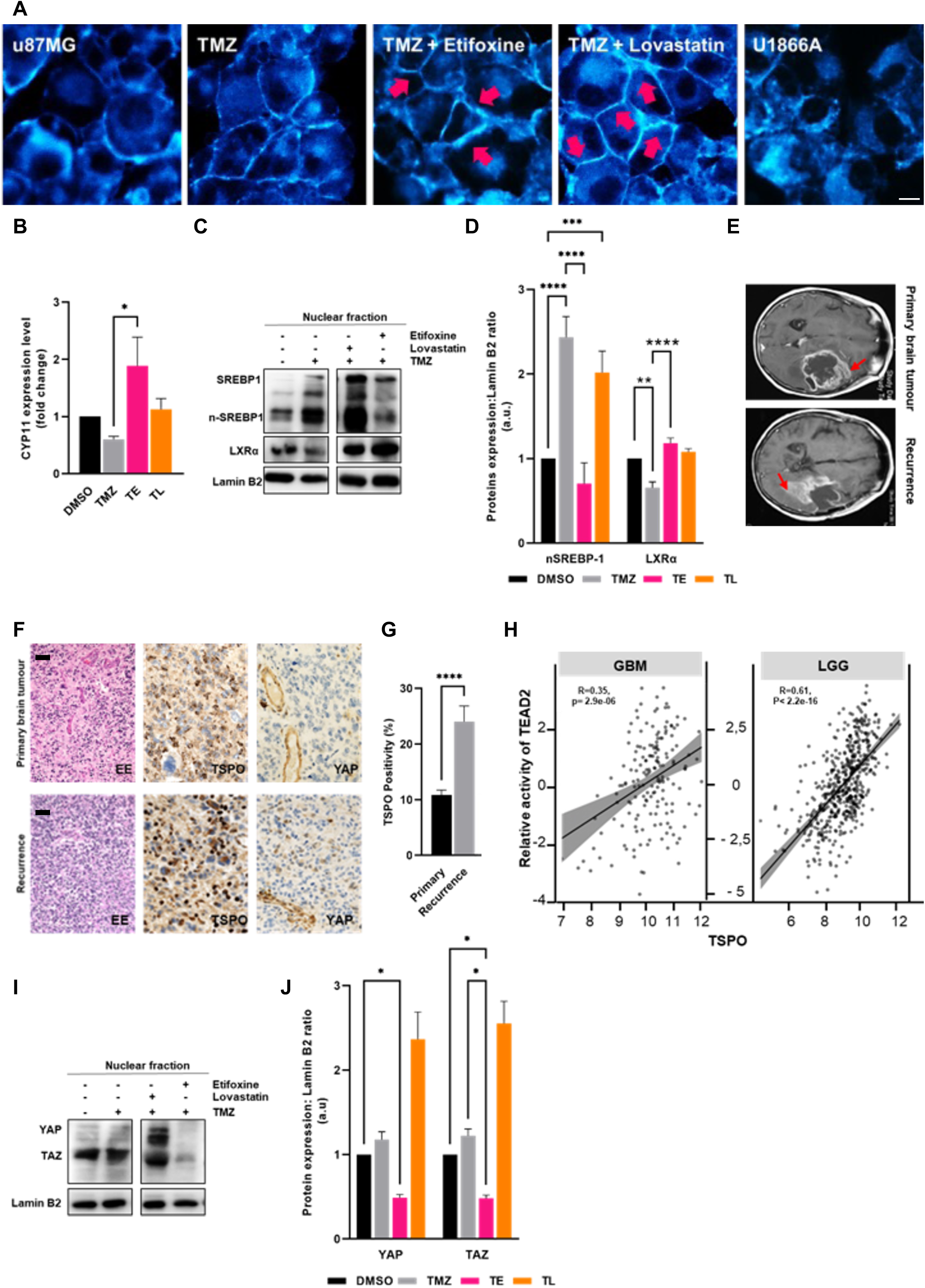
TMZ exploits intracellular cholesterol metabolism and compartmentalization. **(A)** Filipin-based staining of cholesterol (bar=20µm) in U87MG cells treated with TMZ (100μM) alone and combined with Etifoxine (30μM, 24h) and Lovastatin (10μM, 24h). The cholesterol inhibitor U1866A (4μM, 24h) was used as positive control. **(B)** qRT-PCR analysis of CYP11 in U87MG cells treated with TMZ (100μM) alone and combined with Etifoxine (30μM) and Lovastatin (10μM) for 24hours. **(C)** Immunoblotting of LXRα and SREBP1 in nuclear fractions of TMZ-treated U87MG (24h-timepoint, 100μM) alone or combined with Lovastatin (10μM) and Etifoxine (30μM). Quantification (normalized on Lamin B2) is reported in **(D)**. **(E)** Axial post-contrast MRI scan of glioblastoma before surgery and at recurrence during chemotherapy. **(F)** Assessment of TSPO expression in a primary and recurrent case of glioblastoma (bar=50µm haematoxylin-stained sections and immunoperoxidase) quantified in **(G)**. **(H)** Pearson correlation between the relative activity of *TEAD2* transcription factor (normalized enrichment score measured by expression of downstream target genes) and *TSPO* gene expression for TCGA GBM (n=160) and LGG (n=514) tumours. **(I)** Immunoblotting of YAP/TAZ, Actin in nuclear fractions of TMZ-treated U87MG (24h-timepoint, 100μM) alone or combined with Lovastatin (10μM) and Etifoxine (30μM). Quantification (normalized on Lamin B2) is reported in (**J**). All data are represented as mean±sem. *p≤0.05; **p≤0.01; ***p≤0.001; ****p≤0.0001

In addition, being TSPO involved in steroid synthesis (Rupprecht et al., 2010) and the accumulation of intracellular cholesterol coupled with a reduction of the liver X receptor (LXR) transcriptional factor (Bovenga et al., 2015), we investigated the expression of the Sterol Regulatory Element-Binding Protein 1 (SREBP1) and LXRs (**Fig.5C-D**). In response to sterol deprivation, SREBP1 controls the transcription of genes encoding enzymes of the MVA pathway (Mullen et al., 2016). Conversely, LXRs protect the cells from cholesterol overload by stimulating cholesterol efflux (Baranowski, 2008). Our results document the modification in the expression of SREBP1 and LXRs in the nuclear fractions of U87MG cells according to treatments with TMZ (100μM) alone or in combination with Lovastatin (10μM) and Etifoxine (30μM). Our analysis unveiled that the SREBP1 and LXRs expressions in the nucleus vary depending on the treatment. Namely, it showed that TMZ and Etifoxine co-treatments increase LXR whereas SREBP1 expression decreased compared to TMZ treatment alone (**Fig.5C-D**).

### TSPO targeting controls compartmentalization and metabolism of Cholesterol which is key to glioblastoma aggressive behaviour

The MVA pathway (Mullen et al., 2016) controls the transcription factors Yes-associated protein (YAP1) and the transcriptional co-activator with PDZ-binding motif TAZ which we assessed given their relevance with the aggressive behaviour of the glioblastoma. To this end we assessed the TSPO and YAP1 expression in primary tumour and recurrence from adult patients with supratentorial, hemispheric glioblastoma who progressed after radiotherapy and during the cycles of TMZ (**Fig.5E**). Immunostaining revealed considerable expression of TSPO in the recurrent tumours (**Fig 5G**). YAP1 expression paralleled TSPO expression in both the primary and recurrent lesions (**Fig. 5F**). To assess the level of Hippo-YAP pathway activation, we investigated the correlation between TSPO expression and the downstream transcriptional factor TEAD2 using RNA data for TCGA samples (GBM and LGG cohorts). The analysis revealed a strong positive correlation in both datasets (**Fig.5H**). In addition, the accumulation of YAP/TAZ proteins in the nucleus of U87MG cells treated with TMZ (100μM) alone or in combination with Etifoxine or Lovastatin was also detected (**Fig.5I, J**). Our data indicated that the co-treatment with TMZ and Etifoxine blunted the levels of both transcriptional factors suggesting that the action of TMZ can be potentiated by Etifoxine treatment of glioblastoma. In conclusion, ADF represents an ideal model to study the effects of compounds that impair cholesterol biosynthesis on glioma stemness because they can form 3D spheroids (**Fig.6B**) (Malindisa et al., 2019), which show enrichment in stem cell markers SOX2, CD133, Nanog and SCD1 (**Fig.6C**). Hence, both 2D monolayer and 3D spheroid formed by ADF cells were treated for 96hours with different concentrations of TMZ alone and in combination with MF-438, an inhibitor of Stearoyl-CoA Desaturase 1 (SCD1) (**Fig.6D; Fig-6F, G**).

**Figure 6.**
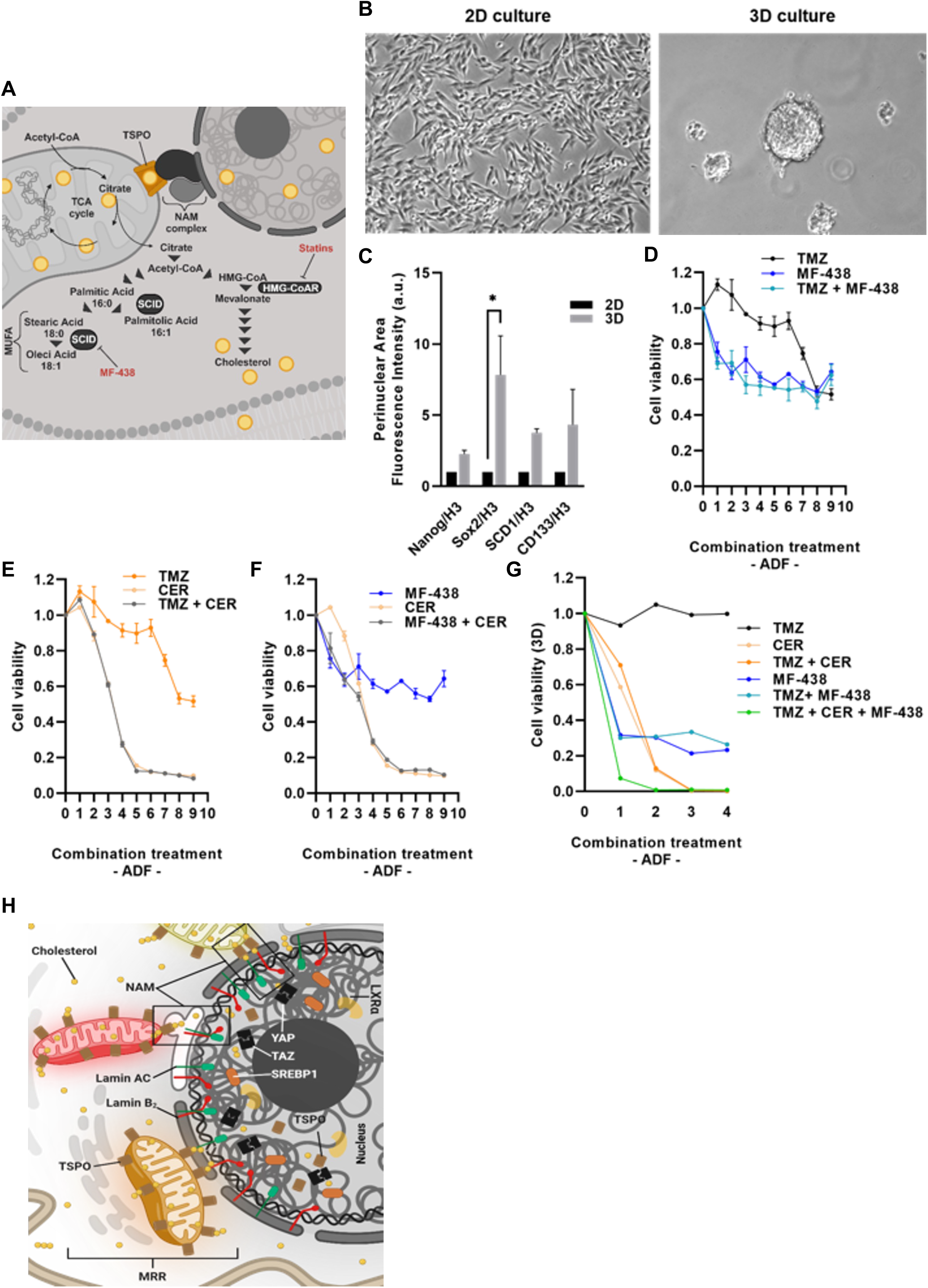
Pharmacological repression of cholesterol promotes TMZ-induced cell death in glioblastoma cells. **(A)** Schematic of lipid metabolism and experimental design for ADF glioblastoma cells treatment. **(B)** Representative images of monolayer (2D culture) and spheroid (3D culture) of ADF cells. **(C)** qRT-PCR analysis of stemness markers (NANOG, SOX2, SCD1, CD133) in 2D versus 3D cultures. Relative expression is shown over H3 mRNA. **(D-F)** Cellular metabolic activity was detected with conventional MTT assay in ADF monolayer cultures (2D) treated with different concentrations of TMZ (0.003, 0.009, 0.3, 0.8, 2.5, 7.4, 22.2, 66.7 and 200μM), Cerivastatin and MF438 (0.008, 0.02, 0.07, 0.2, 0.6, 1.9, 5.6, 16.7 and 50μM), alone or in combination, for 96hours. (**G**) Cell viability was detected via CellTiter Glo in ADF 3D cells treated with TMZ (0.02, 0.14, 3.7 and 33.3μM), Cerivastatin and MF438 (0.008, 0.07, 1.9 and 16.7μM), alone or in combination, for 96hours. **(H)** Working model depicting the role of NAM in the execution of TMZ resistance mechanisms in glioblastoma cells.

The latter is the rate-limiting enzyme for the synthesis of monounsaturated fatty acids and thereby an obvious target to corroborate the role of lipids in the metabolism of glioblastoma cells (**Fig.6A**). Both treatment protocols inhibited the mitochondrial metabolic activity regardless of the 2D or 3D methods of cell growth (**Fig.6D, G**).

Cerivastatin, an inhibitor of the rate-limiting enzyme of cholesterol biosynthesis which targets the 3-hydroxy-3-methylglutaryl coenzyme A (HMG CoA) reductase (**Fig.6A**), was also enrolled in the analysis leading to a greater reduction of cell viability (**Fig.6E-G**). These data confirm that the preservation of cholesterol biosynthesis is pivotal to the proliferation of glioblastoma cells as recently proposed (W. Y. Kim, 2019; Tracz-Gaszewska & Dobrzyn, 2019) and lead to the proposed working model for the resistance mechanism here outlined (**Fig. 6H**).

## Discussion

TSPO regulates the mitochondrial retro-communication with the nucleus to sustain growth and evasion of chemically induced demise (Desai et al., 2020; Strobbe et al., 2021). Such a role is retained in glioblastoma cells exposed to TMZ.

Evolutionary conserved, this 18kDa molecule is embedded within the outer mitochondrial membrane (Beckers et al., 2018) and involved in several cellular processes (Zinnhardt et al., 2021). Over the years TSPO has emerged as a reliable biomarker for the imaging of the pathological brain as during neuroinflammatory and neurodegenerative conditions (Beckers et al., 2018) its expression readily increases in both microglia and macrophages (Betlazar et al., 2018; Wright et al., 2020). TSPO has accumulated in neoplastic cells of several tumours including gliomas in which its increase correlates with the advanced grades of the malignancy (Ammer et al., 2020; Miettinen et al., 1995; Vlodavsky & Soustiel, 2007) whereby it occurs in glioma cells rather than glioma-associated microglia and macrophages (Zinnhardt et al., 2021). Therefore, TSPO ligands have been tested as in vivo tracers of cancer cells and tools to deliver or enhance anti-cancer therapy (Bode et al., 2012). However, the functional role of this potential target in these cells is not yet elucidated. Recently we learned that TSPO partakes in regulating glioblastoma cells’ metabolism (Fu et al., 2020) as its ablation leads to a shift toward glycolysis (Fu et al., 2020). Our *de novo* analysis run by mining data within the Cancer Genome Atlas (TCGA) shows that TSPO upregulation is not beneficial for glioblastoma patients whose survival probability is reduced when the protein is overexpressed (**Fig. 1J**). This is indicative of a threshold level after which TSPO excess principates the malignancy and here we show the alkylating agent TMZ promotes such a negative overexpression which impacts mitochondrial function (**Suppl. Fig. 1,2**). The accumulation of TSPO leads to an increase in oxidative phosphorylation (**Suppl. Fig.1D, E**) but not to an upregulation of the antioxidant mechanisms causing an increase in redox stress (**Suppl. Fig. 1C**) which contributes to the growth (Hayes et al., 2020).

Stabilization of TSPO results in an increased population of mitochondria that resist autophagy (Gatliff & Campanella, 2015; Lawson et al., 2007) (**Suppl. Fig.1D-G; Suppl.Fig2A-D**) despite the prompt formation of autophagosomes following the chemical challenge with TMZ (Filippi-Chiela et al., 2015) (**Suppl. Fig.1I, J**). This aligns well with previous findings for which inhibition of the mitochondria-targeted autophagy (mitophagy) is critical for the pro-survival potential of cells (Desai et al., 2020; S. Kim & Koh, 2017; Yang & Kim, 2019) due to the increased relay with the nucleus (Strobbe et al., 2021). Repressed mitophagy leads to greater mitochondrial content and higher redox stress (Gatliff et al., 2017; Papadopoulos et al., 2006) which initiates and sustains communication with the nucleus via the retrograde pathway (Desai et al., 2020). TMZ exploits this adaptive mechanism by redistributing stressed mitochondria on the nucleus within the range to form contacts (**Fig.2C-I**).

We have described the pro-survival role of NAM, which is now reported in other model systems (Campanella & Kannan, 2024), to be mediated by redistribution of the unmetabolized cholesterol which stations on the nucleus (Ammer et al., 2020) both increasing biosynthesis and accumulation of the lipid (Huang et al., 2020).

We show this to be well recapitulated in TMZ-treated glioblastoma cells in which cholesterol plays a part in aggressive behaviour and chemoresistance (Pirmoradi et al., 2019). TMZ positively regulates the nuclear localization of the transcriptional factor SREBP-1 (**Fig.5C-D**) establishing a positive feed-forward mechanism in which TSPO is required. The TSPO ligand Etifoxine can counteract this adaptive response potentiating the efficacy of the chemotherapeutic agent TMZ (**Fig. 3B**; **Fig. 4A-D**) and the greater the level of TSPO the bigger the co-adjuvant effect of the chemical (**Fig. 3 C, D**).

Etifoxine has been pharmacologically characterized for its allosteric regulation of the GABA receptor signalling and therefore used in clinical practice as an anxiolytic (Strobbe & Campanella, 2018). By upscaling TSPO activity, Etifoxine boosts mitochondrial cholesterol metabolism (Strobbe & Campanella, 2018) clearing the excess in glioblastoma cells which it restates susceptibility to TMZ by removing the excess cholesterol from the nucleus (**Fig.5A-B**). In this a positive regulation of mitophagy by the ligand seems to be a factor as the overall reduction of the mitochondrial network occupancy of the cytosol suggests (**Fig.5E, F**) (**Suppl. Fig.4**).

Etifoxine (Strobbe & Campanella, 2018) does not modify the TSPO expression level in the nucleus hence the physical repositioning of the mitochondria (**Suppl. Fig 4C, D**), but it is still capable of repressing the inter-organelles communication (**Fig.3C, D; Suppl.Fig6A, B**) and the accumulation of transcription factors in the nucleus to protect from cholesterol overload (**Fig.5C, D**). The negative regulation of SREBP-1 isn’t the sole transcriptional pathway controlled by Etifoxine when combined with the TMZ treatment. The nuclear content of the co-transcriptional regulator YAP/TAZ is repressed too (**Fig. 5I, J**). YAP and TAZ are potent co-regulators of stem cell pluripotency which seems favoured by the stabilization of TSPO on the mitochondria. YAP/TAZ integrates genetic and microenvironmental inputs for the amplification, renewal, and proliferation of glioblastoma cells (Castellan et al., 2021).

Previous pieces of evidence have emerged on the chemo-preventive and therapeutic effect of statins in combination with anticancer drugs (Huang et al., 2020).

Here we show that Lovastatin still confers the glioblastoma cells greater susceptibility to TMZ-induced cell death (**Fig. 3B**). Equal effect is recorded with the inhibitors of monounsaturated fatty acids (MUFAs) synthesis which holds the ability to co-adjuvant the TMZ-mediated death in both 2D and 3D glioblastoma cell cultures (**Fig.6 E, G**). Lovastatin is also capable of repressing the nuclear accumulation of the DNA repair enzyme MGMT (O[6]-methylguanine-DNA methyltransferase) by promoting its methylation which is far more affected by Etiffoxine (**Fig.4 E, G**).

We cannot exclude those other signalling mechanisms linked to the MRR that are triggered by TMZ (e.g., Ca^2+^) but we propose that cholesterol has a prominent role in the transduction of mitochondrial inputs to the nucleus in glioblastoma cells. Based on this evidence, TSPO emerges as a potentially valuable target for promoting cholesterol efflux via mitochondria to ameliorate resistance to programmed death. The clarification of the molecules which dictate the tethering of mitochondria with the nucleus will shed light on the mechanistic aspects offering novel avenues to regulate the inter-organelles interplay and approaches to counteract the chemoresistance against TMZ.

## Acknowledgements

The activities in MC labs which led to these findings have been sustained by the following funding bodies we wholeheartedly thank: the European Research Council Consolidator Grant COG2018-819600_FIRM, the LAM-Bighi Grant, the FIRB grant RBFR13P392, the Italian Health Ministry grant IFO14/01/R/52, the Italian Association for Cancer Research grant MFAG21903, Fondation ARC pour la Recherche sur le Cancer ARCLEADER2022020004901 and the Umberto Veronesi Foundation via a persona fellowship to D.S.

We would also like to acknowledge the funding sources that have supported R.M.’s work, including the Italian Association for Cancer Research grant IG24451, Lazio Innova grant A0375-2020-36657, Fondo di Ateneo grant B86C19001510005/RG11916B6F1EE788, and PRIN grant prot.2017HWTP2K.

We would like to express our sincere gratitude to Dr. Del Bufalo (Institute Regina Elena, Rome, IT) and Prof. Barella (University of Rome TorVergata) for generously providing us with the ADF cells and the T98G and A172MG respectively. Thanks go to Dr. Claudia Nardini for her support in the methylation analysis (Department of Pediatric Hematology/Oncology and Cell and Gene Therapy, IRCCS Ospedale Pediatric Bambino Gesù, Rome, IT). Thanks go to Ms Sharmitha Rajendrakumar (Gustave Roussy, FR) for support.

## Author’s Statement

M.C. conceived, designed, and coordinated the project together with D.S. D.S. has performed almost all the experiments, relative analyses, and figures generation. Autophagy experiments were designed by G.L., D.S., E.C.F.C., and M.C. Cell death experiments were designed by R.M. and M.C. Data collection and assembly were performed by D.S., K.B., F.D., C.D.V., I.J.B., F.K., L.S.L., and M.B. G.D. and D.F performed and analysed Immunohistochemical data. L.F. run TEM work. K.B. and E.R. attained and analysed the confocal images. P.I.D. performed surgical isolation of tissues in Figure 7, while I.L. and F.R. performed IHC and analysis. I.N. and A.I. have mined the data on the databases. Manuscript writing was done by D.S., G.L., F.R., and M.C. E.M. and L.P. performed and analysed *MGMT* promoter methylation experiments and contributed to the revision of the manuscript. All authors approved the final manuscript and are therefore accountable for their parts of the study.

