## Supplementary Figures for "Mitochondrial sites of contact with the nucleus aid in chemotherapy evasion of glioblastoma cells"

#### Supplementary Figure 1

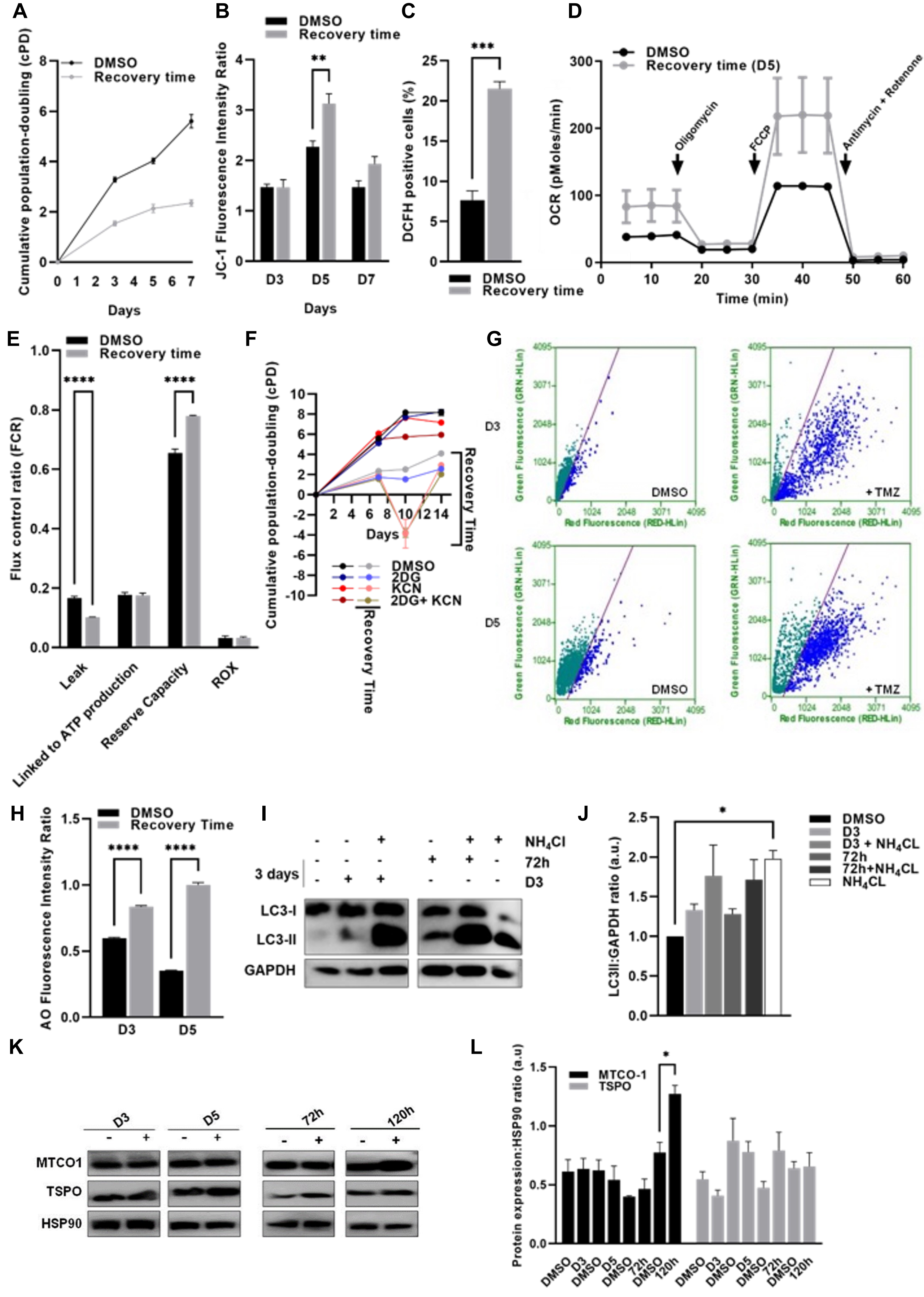

### Supplementary Figure 2

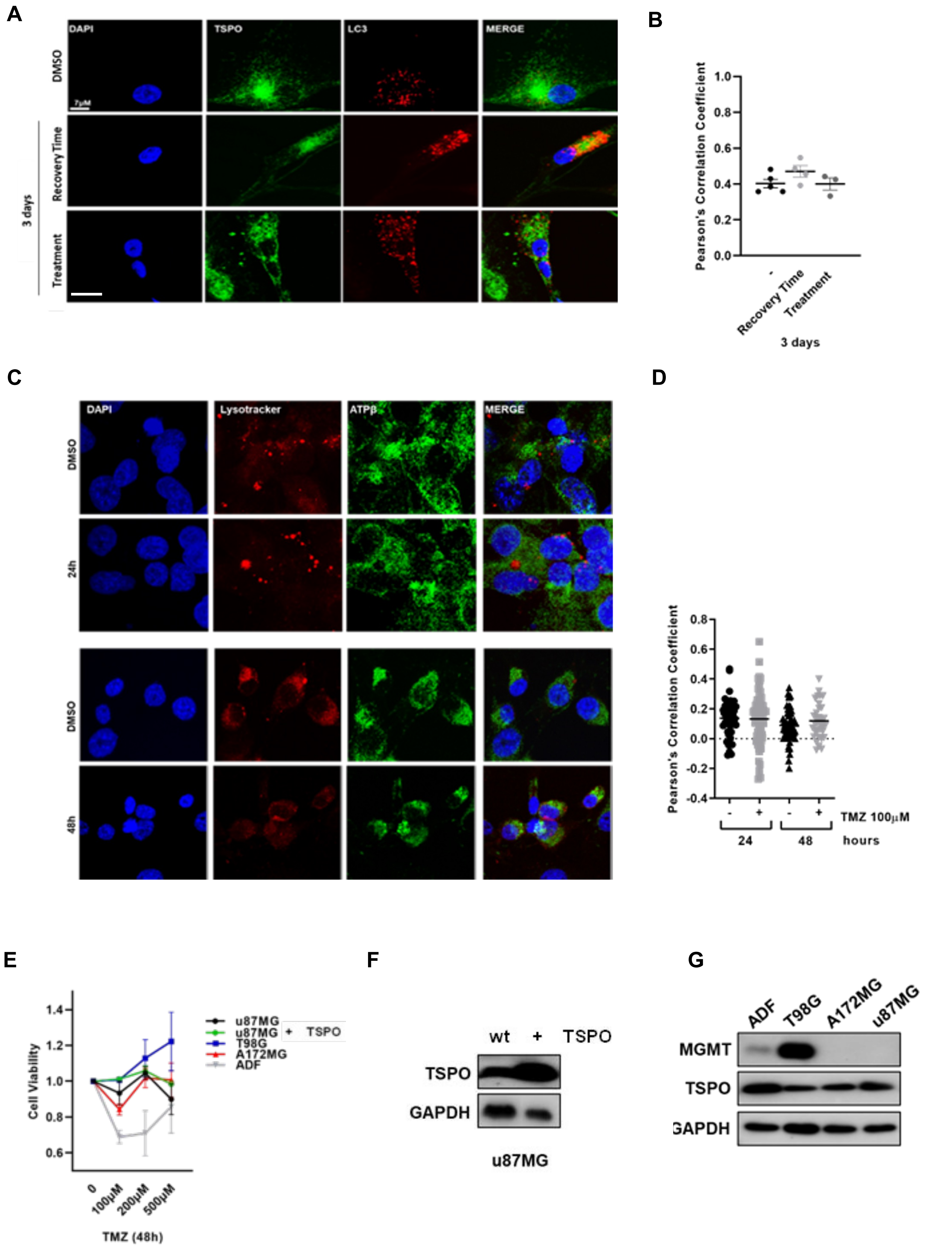

#### Supplementary Figure 3

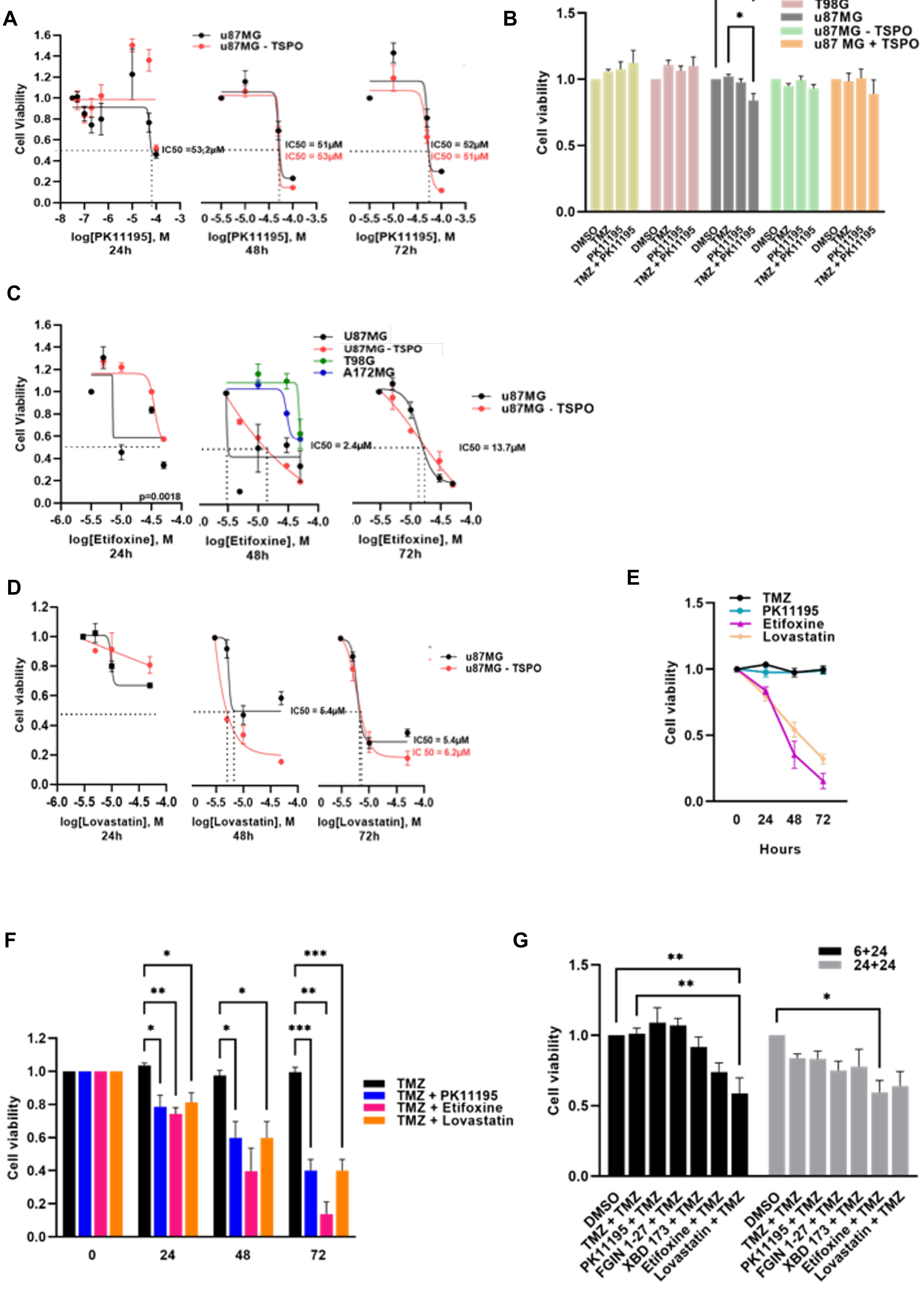

#### Supplementary Figure 4

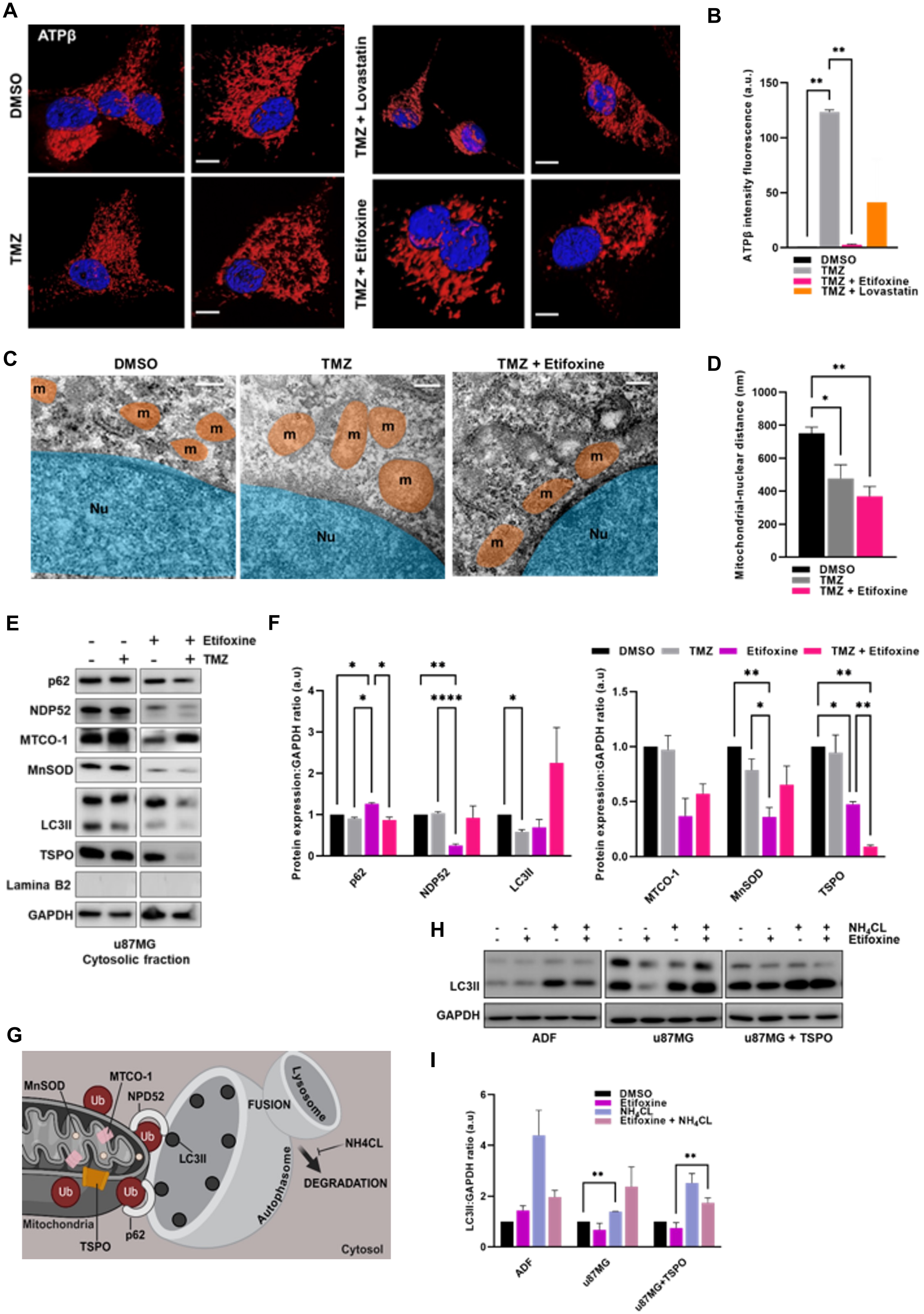

**Supplementary Figure 5**

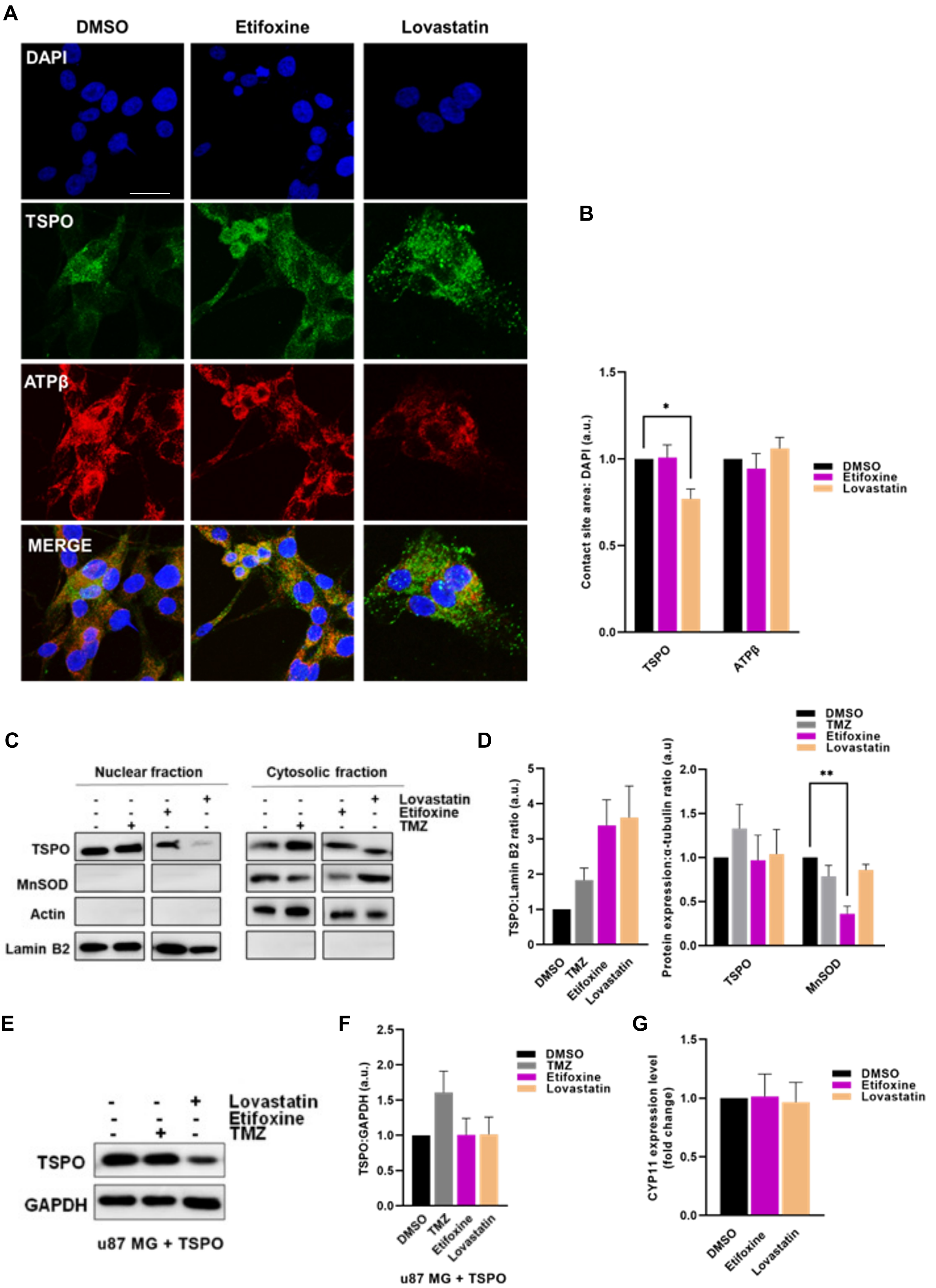

#### Supplementary Figure 6

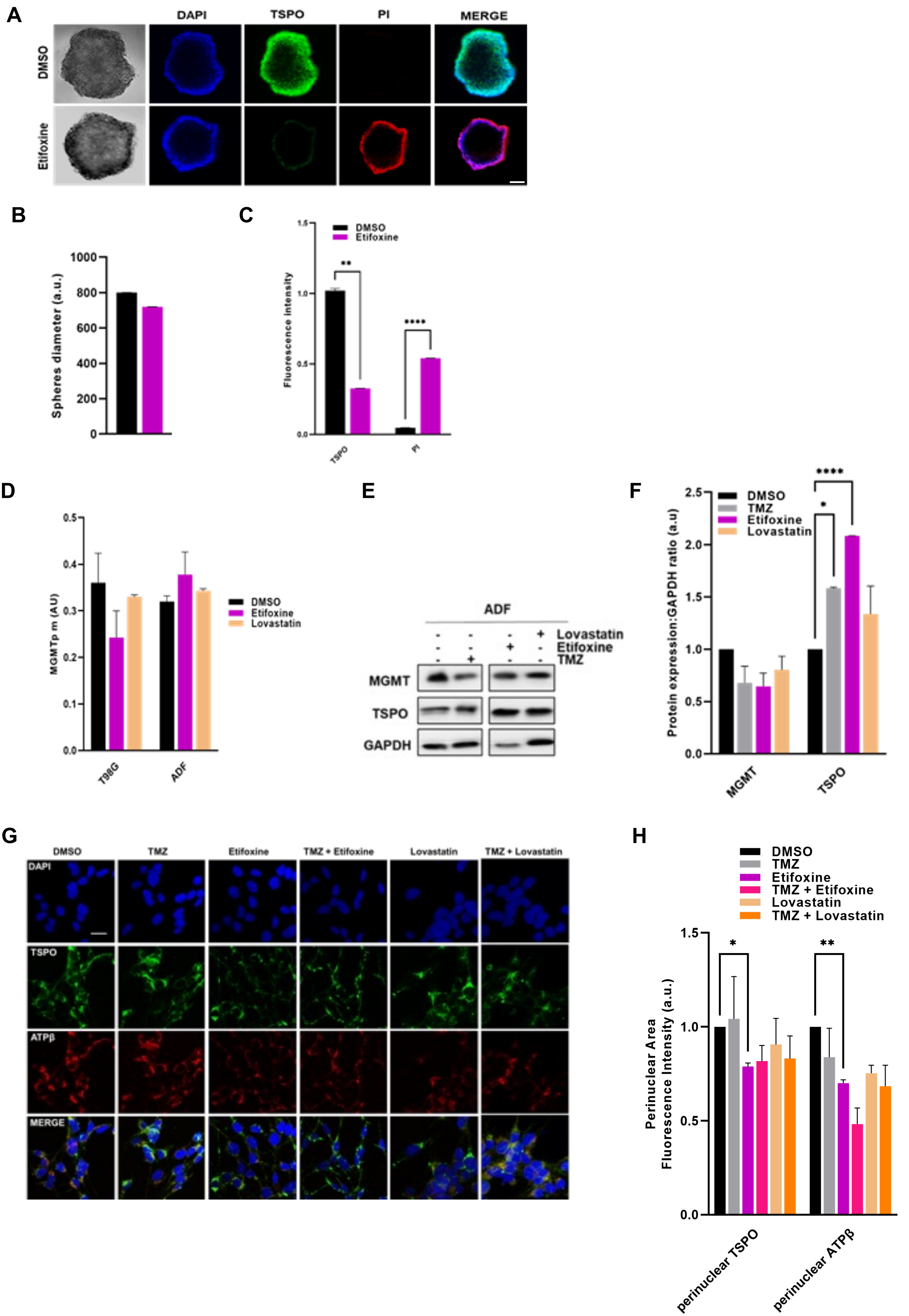

#### Legends to Supplementary Figures

**Supplementary Figure 1. TMZ mediated regulation of mitochondrial homeostasis. (A).** Cumulative Population Doubling (cPD) in u87MG control and TMZ-treated cells on D3, D5 and D7. The u87MG cells were treated with TMZ (100 µM) and then kept in Drug-Free Medium for 3 (D3), 5 (D5), or 7 (D7) days (Recovery Time). **(B)** Mitochondrial membrane potential ($\Delta\Psi_m$) was assessed on D3 in u87MG cells. **(C)** U87MG cellular oxidative stress on D5 was assessed via DCFH-DA dye. **(D)** Oxygraphy assay to measure OCR (oxygen consumption rate) in u87MG cells on D5. **(E)** FCR (fluorescence recovery after photobleaching) analysis on D5. **(F)** The cPD in u87MG control and TMZ-treated cells at several time points in presence of KCN (137 µM and 40 µM respectively for u87MG and u87MG + TMZ) and 2-DG (1.6mM). To assess cell number control cells and cells previously treated with TMZ (100 µM, 3h) were treated with KCN and 2DG on D5. The CPD was determined on D7. **(G)** Flow cytometry analysis of acridine orange (AO) staining in u87MG control and TMZ-treated cells at two timepoints. **(H)** Histogram representation of AO on D3 and D5 **(I)** Immunoblotting analysis of LC3B-II in u87MG cell lysates at D3 and 72 hours of TMZ exposure combined with 24 hours NH4CL (20nM). The graph in panel **(J)** shows the relative densitometry of LC3B-II normalized to HSP90 (a loading control) at different timepoints of TMZ treatment with or without NH4CL. **(K)** Representative western blotting analysis of MTCO-1 and TSPO in protein extracts isolated from cells exposed to both D3/D5 and 72/120 hours TMZ treatment. Densitometry analysis of MTCO-1 and TSPO normalized to HSP90,are quantified in (L).

**Supplementary Figure 2. TMZ mediated patterns of autophagosomes and lysosomes in glioblastoma cells. (A)** Representative Immunofluorescence (IF) of TSPO (green) and LC3 (red) after TMZ treatment (100µM; D3 and 72 hours). DAPI was used to stain nuclei. Scare bar is 7µm. Co-localization analysis was performed by ImageJ plugin Jacop and reported in **(B)**. **(C)** Representative IF of u87MG cells exposed to TMZ (100µM) for 24 (top row) and 48 hours (bottom row) labelled with ATPβ (green) and Lysotracker red. DAPI was used to stain nuclei. Co-localization analysis was performed by ImageJ plugin Jacop and reported in **(D)**. **(E)** Dose-response curve evaluated via Crystal Violet assay in u87MG, ADF, T98G and A172MG cell lines treated with TMZ (100, 200 and 500µM) for 48 hours. **(F-G)** Representative western blotting analysis of TSPO and MGMT in protein extracts isolated from u87MG + TSPO cells and different human glioblastoma cell lines, respectively.

**Supplementary Figure 3. TSPO validation as target to control susceptibility of glioblastoma cells to TMZ.** Cell viability assay was evaluated via crystal violet in both u87MG and u87MG - TSPO cells treated with PK11195 (50,100,200 and 500mM; 10, 50 and 100µM) (**A**), Etifoxine (5, 10, 30 and 50µM) **(C)** and Lovastatin (5, 10, 50µM) **(D)** at different concentrations (from 0 to 100µM) for 24-72 hours. **(B).** Cell viability assay was also evaluated via crystal violet in u87MG, u87MG – TSPO and u87MG + TSPO, A172MG and T98G cells treated with TMZ (100µM) alone and combined with PK11195 (200nM) for 24 hours. Effect of TSPO ligands alone (PK11195 (200nM), Etifoxine (30µM) and Lovastatin (10µM) **(E)** and combined with TMZ (100µM) **(E-F)** were also detected via crystal violet assay in u87MG cells after 24-72 hours treatment. (G). Cell viability was evaluated in u87MG cells treated with TSPO ligands (PK11195 200nM; FGIN 1-25 200nM; XBD 173 30µM; Etifoxine 30µM; Lovastatin 10µM) for 6 and 24 hours prior to being treated with TMZ (100µM) for 24 hours

Data are shown as the mean ± SEM. ****P<0.0001, ***P<0.001, **P< 0.01 and *P<0.05.

**Supplementary Figure 4. Etifoxine counteracts TSPO mediated inhibition of mitophagy.** **(A)** Confocal images (bar=10μm) with isosurfaces for red channel (Imaris, Bitpalne) showing ATPβ fluorescence intensity after TMZ treatment (100μM) combined with Lovastatin (10μM) and Etifoxine (30μM) for 24 hours. ATPβ expression was quantified in **(B)**. **(C)** TEM micrographs of u87MG untreated and treated cells with TMZ 100μ, 24 hours combined with Etifoxine (30μM, 24hours) to confirm the redistribution of physical interaction between mitochondria (M; orange) and the nucleus (N; blue) as represented in **(D)**. **(E)** Representative western blotting of autophagic proteins in cytosol of u87MG after 24h of incubations with TMZ (100μM) alone and in combination with Etifoxine (30μM). Protein extracts were analysed to detect p62 (Abcam) NPD52(Proteintech), MTCO1, LC3 and nuclear Lamin as indicated. Densitometry analysis protein levels normalized over GAPDH is also reported in **(F)**. **(G).** Schematic mechanism of mitophagy. **(H)** Immunoblotting analysis of LC3B-II in u87MG, u87MG + TSPO and ADF cell lysates following exposure to Etifoxine (30μM) for 24 hours alone and in combination with TMZ (100μM), and in the presence of NH4CL for 24hours. The graphs in panels (**H)** show the relative densitometry of LC3B-II normalized to GAPDH.

**Supplementary Figure 5.** **(A)** Isosurfaces derived from confocal images of immunocytochemical analyses of TSPO (green) and ATPβ (red) at 24h-time point of Lovastatin (10μM) and Etifoxine (30μM) treatment (bar=100μ). **(B)** Quantification of the mito-nuclear contacts. **(C)** Immunoblotting of TSPO, MnSOD, Actin and Lamin B2 in nuclear and cytosolic fractions of u87MG cells treated with Lovastatin(10μM) or Etifoxine (30μM) for 24 hours. **(D)** Reports densitometry analysis of TSPO and MnSOD levels. **(E)** Representative western blotting of TSPO expression in total lysates of U87MG + TSPO after 24 hours of incubations with Lovastatin (10μM) and Etifoxine (30μM). Densitometry analysis of protein levels normalized over GAPDH is reported in panel **(F).** **(G).** qRT- PCR analysis of CYP11 in U87MG cells treated with  with Etifoxine (30μM) and Lovastatin (10μM) for 24 hours.

**Supplementary Figure 6.** (**A**) Representative images of U87MG spheroids stained with Dapi, PI and antibody for TSPO and exposed to 30 μM Etifoxine. Histograms reporting quantification of spheroids diameter in the condition of analysis quantified over control (**B**) and relative changes in fluorescent intensity (**C**). **(D)** Percentage of MGMT promoter methylation (MGMTp meth%) in T98G and ADF cell lines. **(E, F)** Immunoblotting of MGMT and TSPO in ADF total lysates treated with Lovastatin (10μM) and Etifoxine (30μM) for 24 hours. The graph in panel **(F)** shows the relative densitometry of MGMT and TSPO normalized to GAPDH. **(G)** ADF isosurfaces derived from confocal images of immunocytochemical analyses of TSPO (green) and ATPβ (red) at 24h-time point of TMZ treatment (100μM)combined with Lovastatin (10μM) and Etifoxine (30μM) (bar=100μ). **(H)** Quantification of the mito-nuclear contacts

All data are represented as mean±sem. *p≤0.05; **p≤0.01; ***p≤0.001

**Material and methods**

**Cell viability.** The cPD was performed in u87MG cells (15.000 cells per well) treated with TMZ 100µM for 3h and then cultured in drug-free medium for 3 (D3) and 5 (D5) day. It was also calculated u87MG control and TMZ-treated cells at several time points in presence of KCN (137 µM and 40 µM respectively for u87MG and u87MG + TMZ) and 2-DG (1.6mM). cPD refers to a number of times a population of cells has doubled in number and calculated as = log (Nf/N1)/log2.

**Measurement mitochondrial membrane potential (ΔΨm).** The u87MG cell lines were plated (15.000 cells per well) 24 hours before treatment. The Δψm was detected by staining cells with 2µM JC1 (Thermo Fisher, T3168) for 20 min at 37°C and 5% CO2. Sample were collected and analysed by GUAVA EasyCyte flow cytometer on D3, D5 and D7.

**Oxidative stress**. The u87MG cell lines were treated with 100µM TMZ for 3h, cultured in drug-free medium for 5 (D5) days, washed with PBS, harvested with trypsin (Gibco, 15090046), suspended in warm medium containing 10µM dichloro-dihydro-fluorescein diacetate (DCFH-DA; Invitrogen, D399) for 20 min at 37°C and analysed by flow cytometry.

**High resolution respirometry system.** Mitochondrial oxygen consumption assays were performed using the high-resolution respirometry system Oxygraph-2 k (Oroboros Instruments, Austria). Cell respiration was measured at the density of 1.5 x 106 for the control cells and 1 x 106 for the treated cells. The assay was made at 37°C in 2mL chambers containing culture medium at stirring rate of 750 rpm. Respiration (R) was measured in the coupled state of physiological respiratory control. We also used oligomycin (2µg/mL, Sigma, 75351), which inhibits the mitochondrial phosphorylation, to determine the non-coupled resting respiration, or LEAK respiration (L). FCCP at the concentration of 1.5 µM (Abcam, ab120081) was used as an optimum uncoupler, providing a measure of the kinetic capacity of the electron transport system (ETS) and helping us to measure the maximal uncoupled respiration (E). Using the ETS capacity as a common basis for normalization of coupling control ratios, the R/E reflects the level of mitochondrial activity relative to the maximal kinetic capacity of the electron transport system. Correspondingly, the L/E ratio reflects the level of LEAK respiration relative to the ETS capacity and provides an estimate of intrinsic uncoupling. Finally, the fraction of respiration used for ATP production was estimated as (R-L)/E. The analyses were performed on D5.

**Assessment of the autophagy levels.** The u87MG cell lines were plated (15.000 cells per well) 24 hours before treatment (3hours, TMZ 100µM). Then, the cells were cultured for 3 (D3) and 5 (D5) days, collected by trypsinization, stained with 2,7µM Acridine Orange (AO, Thermo Fisher, A1301) for 15 min at room temperature and analysed on a GUAVA EasyCyte flow cytometer using InCyte 2.6 software (Guava Technologies), as previously described (doi:10.1242/jcs.195057). To further investigate the role of autophagy, the u87MG cells, cultured for 3 days (D3) and 72 hours, were also treated with NH4CL (20nM) for the final 24 hours. Western blotting analysis was conducted using an LC3 (Novus Biologicals, NB100-2220).

**Immunofluorescence microscopy.** Regarding LysoTracker™ Red DND-99 (Invitrogen, L7528), the u87MG cells were preincubated with the dye for 30 min before being washed with PBS and fixed with 4% PAF. The co-localization was analysed by linear regression and calculated by Pearson correlation coefficient (Image J software).
