## Supplementary Methods for "Mitochondrial sites of contact with the nucleus aid in chemotherapy evasion of glioblastoma cells"

### Cell culture

GBM cell lines u87MG, T98G and A172MG were grown in Dulbecco's Modified Eagle Medium (DMEM, Gibco) whereas ADF was cultured in RPMI 1640 (Corning) supplemented with 10% FBS (Lonza), 100U/mL penicillin and 100mg/mL streptomycin (Life Technologies) at 37°C and 5% $CO_2$.

### Treatment of cell

Applied cell lines were treated with Temozolomide (Sigma, T2577), PK11195 (Enzo Life Technologies, BML-CM 118), Etifoxine (Sigma, SML 0272), Lovastatin (Enzo, BML-G226), Cerivastatin (Sigma, SML0005) and MF-438 (Sigma, 569406) (was kindly provided by Ziga Jakopin) at different concentrations for different times.

### Molecular modulation of TSPO

The downregulation of TSPO in u87MG cell line (u87MG - TSPO) was achieved using the pGIPZ shRNA vector: clone ID V3LH_331646, target sequence: 5′-TGAGTGTGGTCGTGAAGGC-3, purchased from Open Biosystems (Huntsville, AL, USA). The plasmid contains the TurboGFP (tGFP) reporter for visual tracking of transduction and expression. Transfection was performed using standard $Ca^{2+}$ phosphate-based protocol as previously demonstrated[1]. After transfection, cells were maintained for 2 weeks in media supplemented with 3µg/ml puromycin (SERVA Electrophoresis GmbH, 33835) for the selection of transfected (GFP-positive) cells. TSPO downregulation in u87MG - TSPO cells was confirmed by western blot analysis.

The over expression of TSPO in u87MG cell line (u87MG + TSPO) was performed using the TSPO encoding plasmid from (Origene RG220107). The plasmid contains the C-terminal MYC/DDK tag, reporter for visual expression. Transfetion was achieved using Lipofectamine2000 as typically provided by the manufacturer. After transfection, the transfected cells were selected using G418 at a concentration 700 µg/µL. Cellular pellet was then subjected to Western Blot analysis to confirm the overexpression of TSPO.

### Cell viability

Cumulative population doubling (cPD) was performed in u87MG cells (15.000 cells per well) treated with TMZ 100µM for 3h and then cultured in drug-free medium for 3 (D3) and 5 (D5) day. It was also calculated u87MG control and TMZ-treated cells at several time points in presence of

KCN (137 μM and 40 μM respectively for u87MG and u87MG + TMZ) and 2-DG (1.6mM). cPD refers to a number of times a population of cells has doubled in number and calculated as = log $(N_f/N_1)/\log_2$. $N_f$ is the final number of cells whereas $N_1$ is the initial number of cells.
Cell viability assay in GBM cell lines was measured using crystal violet solution (Sigma, HT90132). The cells were treated with TMZ at different concentrations, in combination with TSPO ligands (PK11195 and Etifoxine) or Lovastatin at different concentrations for different time points. A curve dose-response was also measured for both TSPO ligands and Lovastatin as showed in Supplementary Figures. After treatment, cellular viability was assessed by crystal violet assay as previously described[2].
Mitochondrial metabolic activity of ADF was detected via 3-(4,5-dimethylthiazol-2-yl)-2,5-diphenyltetrazolium bromide (MTT, Invitrogen, M6494) assay[3]. Cell Titer Glo 3D (Promega, G9683) was used to evaluate cellular viability of ADF in 3D cultures as previously demonstrated[4].

**Sphere formation assay**

Sphere propagation assays were performed as previously described[5]. Briefly, single-cell preparation (1000 cells/well) of ADF cell lines were suspended in an appropriate amount of sphere-forming medium: serum-free DMEM/F12 (Gibco) supplemented with basic Fibroblast Growth Factor (bFGF, Sigma Aldrich, F0291), Epidermal Growth Factor (EGF, Sigma Aldrich, E4127), insulin (Sigma Aldrich, 9011-M), glucose (Sigma Aldrich, G7021), heparin (Sigma, 2106), B27 (Gibco, 17504001) and plated into a 96-well ultra-low adherent plate (Costar) to form spheres. The documentation of images and evaluation of sphere-forming efficiency were performed on 96h.

**Assessment of the autophagy levels**

U87MG cell lines were plated (15.000 cells per well) 24h before treatment (3h, TMZ 100μM). Then, the cells were cultured in drug-free medium for 3 (D3) and 5 (D5) days, collected by trypsinization, stained with 2,7μM Acridine Orange (AO, Thermo Fisher, A1301) for 15 min at room temperature and analysed on a GUAVA EasyCyte flow cytometer using InCyte 2.6 software (Guava Technologies), as previously described[6].

**Subcellular fractioning and Western blot analysis**

After treatment, GBM cells were washed in Phosphate Buffer Saline (PBS, Gibco), collected by gentle scraping, and incubated in ice-cold RIPA buffer (50mM Tris–HCl, 150mM NaCl, 1% (v/v) TritonX100, pH 8.0) containing protease and the phosphatase inhibitor for 30 min on ice. Total lysates were centrifuged at 13,000*g* at 4°C for 20 min. Subcellular fractionation was performed with Rapid Efficient And Practical (REAP) method[7]. GBM cells were incubated in Reap Lysis Buffer (PBS 1X and 0,1% Nonidet P-40 Substitute) for 3 min on ice. Lysates were centrifuged at 2200g for 5 min at 4°C. The pellet corresponds to nuclear fraction was washed, centrifuged 3 times at 2200rpm for 5 min at 4°C and sonicated 3 times for 5 sec, amplitude 80. Protein concentration was estimated using the Bradford reagent (Biorad, 5000006), and 20-40µg of total proteins was mixed with Laemmli sample buffer (Biorad) and boiled at 95°C for 2 min. Proteins were resolved by Sodium Dodecyl Sulfate Polyacrilamide gel (SDS–PAGE) and transferred to PVDF (Merk Millipore) membranes. Membranes were blocked in 5% Bovine Serum Albumine (BSA) (Sigma, A7030) or 5% non-fat dry milk (Applichem, A0830) in 1×Tris buffer saline (TBS) (25 mMTris, 0.15 M NaCl) containing 0.05% Tween-20 (Sigma, P9416) (TBST, pH 7.5) for 1h, probed with appropriate diluted primary (OXPHOS (Abcam, ab110412), LC3 (Novus Biologicals, NB100-2220), GAPDH (Abcam, ab9485), MTCO1 (Abcam, ab203912), TSPO (Abcam, ab118913), OPA1 (BD, 612607), MFN1 (Santa Cruz. Sc-166644), HSP90 (Santa Cruz, sc-69703) Lamin A/C (Cell signalling, 4777), Lamin B2 (Thermofisher Scientific, 33-2100), ATPβ (Abcam, ab14730), Actin (Sigma-Aldrich, A2066), MnSOD (Millipore, 06-984), SREBP-1 (Santa Cruz, sc-13551), YAP (Santa Cruz, sc-101199), LXR (Abcam, ab3585) and CASP 3 cleaved (Cell signalling, 9661) overnight at 4°C washed in TBST (3×10 min at RT) and then probed with the corresponding peroxidase-conjugated secondary antibody (Biorad) for 1h at RT.

After further washing in TBST (3x10min at RT), proteins were detected using an ECL detection kit (Merk Millipore) via Amersham Imager 600. Immunoreactive bands were analysed by performing densitometry analysis with ImageJ software (NIH, Bethesda, MD, United States).

**High resolution respirometry system**

Mitochondrial oxygen consumption assays were performed using the high-resolution respirometry system Oxygraph-2 k (Oroboros Instruments, Austria). Cell respiration was measured at the density of 1.5 x $10^6$ for the control cells and 1 x $10^6$ for the treated cells. The assay was made at 37°C in 2mL chambers containing culture medium at stirring rate of 750 rpm. Respiration (R) was measured in the coupled state of physiological respiratory control. We also used oligomycin (2µg/mL, Sigma, 75351), which inhibits the mitochondrial phosphorylation, to determine the non-coupled resting respiration, or LEAK respiration (L). FCCP at the concentration of 1.5 µM (Abcam, ab120081) was used as an optimum uncoupler, providing a measure of the kinetic capacity of the electron transport system (ETS) and helping us to measure the maximal uncoupled respiration (E). Using the ETS capacity as a common basis for normalization of coupling control ratios, the

R/E reflects the level of mitochondrial activity relative to the maximal kinetic capacity of the electron transport system. Correspondingly, the L/E ratio reflects the level of LEAK respiration relative to the ETS capacity and provides an estimate of intrinsic uncoupling. Finally, the fraction of respiration used for ATP production was estimated as (R-L)/E. The analyses were performed on D5.

**Measurement of mitochondrial mass and mitochondrial membrane potential ($\Delta\Psi_m$)**

U87MG cell lines were plated (15.000 cells per well) 24h before treatment, which consisted of 3h with TMZ 100µM. Then, the cells were cultured in drug-free medium for 3 (D3), 5 (D5) and 7 (D7) days. The mitochondrial mass was assessed upon staining the cells with 200nM Mitotracker Green FM (Thermo Fisher, M7514) for 20 min at 37°C and 5% $CO_2$. The mitochondrial membrane potential ($\Delta\psi_m$) was detected by staining cells with 2µM JC1 (Thermo Fisher, T3168) for 20 min at 37°C and 5% $CO_2$. Sample were collected and analysed by GUAVA EasyCyte flow cytometer on D3, D5 and D7.

**Oxidative stress**

U87MG cell lines were treated with 100µM TMZ for 3h (acute treatment), cultured in drug-free medium for 5 (D5) days, washed with PBS, harvested with trypsin (Gibco, 15090046), suspended in warm medium containing 10µM dichloro-dihydro-fluorescein diacetate (DCFH-DA; Invitrogen, D399) for 20 min at 37°C and analysed by flow cytometry.

**RNA extraction, reverse transcription, and quantitative real-time PCR**

U87MG cells were lysed using GENEAID - Total RNA Mini Kit (Geneaid Biotech Ltd, RBD100) and total RNA was extracted using TAKARA - PrimeScript RT Reagent Kit (Takara, RR037A) according to the manufacturer's instructions. In all, 1µg of RNA was retro-transcribed using SensiFAST™ cDNA Synthesis Kit (Bioline) and used in quantitative RT-PCR (qPCR) experiment, using SensiFAST™ SYBR Hi-ROX Kit (Bioline) following manufacturer's instructions. Thermocycling consisted of an initial polymerase activation step at 98 °C for 5 min, and amplification was performed with 35 cycles of 95 °C for 15 s, 68 °C for 10 s, and 72 °C for 20 s with data acquisition at this stage and the reaction finished by the built-in melt curve. The relative amounts of mRNA were calculated by using the comparative Ct method. Primers for *GAPDH*: 5'-TCATGGGTGTGAACCATGAGAA-3' (forward) and 5'-GGCATGGACTGTGGTCATGAG-3' (reverse); *TSPO*: 5'-TCTTTGGTGCCCGACAAATGG-3' (forward) and 5'-

TCAACTACTGCGTATGGCGGGACAACC-3’ (reverse); *CYP11*: 5’-CCGTGACCCTGCAGAGATAT-3’ (forward) and 5’- TGGTCATCTCTAGCTCAGCG-3’ (reverse); *SREBP1*: 5’-CGGAACCATCTTGGCAACA-3’ (forward) and 5’-GCCGGTTGATAGGCAGCTT-3’ (reverse)

Total RNA of ADF were extracted and isolated by TRIzol (Thermo Fisher, 15596018) following the manufacturer’s instructions[8]. Quantitative RT-PCR was performed as previously described[5,9]. Gene expression was normalised to H3 mRNA content.

**Immunohistochemistry**

Following PBS washing, primary human GBM sections were incubated at 80°C for 30m in citrate buffer: 10mM Sodium Citrate dihydrate (Sigma, W302600), 0.05% Tween-20 (Sigma, P9416), pH 6.0. Washes were flowingly carried out in PBS-T (0.05% Tween-20 in PBS). Permeabilization was carried out for 15 min at 4°C in 20mM HEPES, 300mM Sucrose (Sigma, S0389), 50mM NaCl, 0.05% Triton-X100 (Sigma, T8787), 3mM $MgCl_2$, 0.05% Sodium Azide (Sigma, S2002). Blocking – 1h at room temperature in 10% horse serum (Thermo Fisher), 1% w/v Bovine Serum Albumin (BSA, Applichem, A1391) in PBS. TSPO (Abcam, ab118913) primary antibody incubation was carried out overnight at 4°C followed by the secondary antibodies anti-rabbit (Invitrogen, A-11012) in blocking solution for an hour at RT, while shaking. Slices were washed 4-5x in PBS-T and 3x in PBS, before a 5 min incubation with 600nM DAPI solution (Applichem, A4099) for nuclei visualization. Excess DAPI was cleared with 3x5 min washes in PBS and slices were then mounted on glass slides (VWR) using a ProLong Gold antifade reagent (Invitrogen, P36930), covered with 22 mm glass coverslips. The slices were then imaged by means of an Olympus Fluoview 1000 Confocal Laser Scanning Microscope using objective 60x oil (N.A. 1,35) and laser 405nm for DAPI fluorescence, 488nm for green fluorescence and 543nm for red fluorescence. All the images were processing with Imaris software (Bitplane, Switzerland) for 3d rendering and isosurfaces and analyzed with ImageJ software for colocalization analysis. All images were analysed by ImageJ software (NIH, Bethesda, MD, United States).

**Immunofluorescence microscopy**

U87MG cells were initially plated on 22 mm glass coverslips at 70% confluency. At treatment completion, primary human GBM cells were washed once with Phosphate Buffered Saline (PBS) and fixed with 4% paraformaldehyde (PFA) following an incubation of 10 minutes. Cells were then washed 3 times for 5 minute each with PBS and permeabilized with a 0.1% Triton-X-100 (Applichem, A4975)/PBS solution for 10-20 minutes. After another round of washes cells were blocked for 1 hour in a solution of PBS containing 3% w/v BSA. Primary antibodies ATPβ (Abcam, ab14730), TSPO (Cell Signalling, 70358), NF-kB (Abcam, ab16502), LC3 (Novus Biologicals, NB100-2220) and Lysotracker red (Invitrogen, L7528) which were then incubated overnight at 4°C in blocking solution. After 3 washes, the secondary antibody was incubated for 1 hour in blocking solution: anti-mouse (Invitrogen, A-11094) and anti-rabbit (Invitrogen, A-11012). After 1 hour at room temperature away from light, the coverslips were washed and mounted onto glass slides using a 4',6'-diamidino-2-phenylindole (DAPI)-containing mounting medium (ab104139). The coverslips were then imaged on an Olympus Fluoview 1000 Confocal Laser Scanning Microscope using objective 60x oil (N.A. 1,35) and laser 405nm for DAPI fluorescence, 488nm for green fluorescence and 543nm for red fluorescence. All the images were processing with Imaris software (Biplane, Switzerland) for 3d rendering and isosurfaces and analyzed with ImageJ software for colocalization analysis.

Regarding LysoTracker™ Red DND-99 (Invitrogen, L7528), u87MG cells were preincubated with the dye for 30 min before being washed with PBS and fixed with 4% PAF.

For detection of cellular cholesterol, we used a cell-based cholesterol assay kit (ab133116; Abcam, Cambridge, MA). Briefly, u87MG cells were fixed with PFA (4%) and stained with filipin III following the manufacturer instructions. Cells were visualized using microscope Zeiss Axio observer 7.

**Statistical analysis**

Statistical analyses were performed using GraphPad Prism 9 software, choosing the most appropriate test. Number of independent experiments and number of replicates used to perform statistics is ≥ three. Unpaired Student's t test and ordinary one or two-way ANOVA followed by Bonferroni Test were used for cellular correlation and multiple comparisons. Post-hoc test was conducted only if F achieved $P<0.05$. Moreover, the co-localization was analysed by linear regression and calculated by Pearson correlation coefficient. Data are presented as means ± standard error of the mean (SEM). Statistical significance was declared at $p \leq 0.05$. The asterisks describe different values levels of statistical significance: ****$P<0.0001$, ***$P<0.001$, **$P< 0.01$ and *$P<0.05$.
